# Predicting evolution in experimental range expansions of an aquatic model system

**DOI:** 10.1101/2022.01.20.477128

**Authors:** Giacomo Zilio, Sascha Krenek, Claire Gougat-Barbera, Emanuel A. Fronhofer, Oliver Kaltz

**Affiliations:** ISEM, University of Montpellier, CNRS, EPHE, IRD, Montpellier, France.; Institute of Hydrobiology, Technische Universität Dresden, Dresden, Germany

**Keywords:** Dispersal evolution, reaction-diffusion model, predictability, eco-evolutionary dynamics, invasions

## Abstract

Predicting range expansion dynamics is a challenge for both fundamental and applied research in conservation and global change biology. However, if ecological and evolutionary processes occur on the same time scale, predictions are challenging to make. Combining experimental evolution and mathematical modelling, we assessed the predictability of independent realisations of range expansions in a laboratory model system, the freshwater protozoan *Paramecium caudatum*. We followed ecological dynamics and evolutionary change in range core and front populations in the experiment. These settings were recreated in a predictive mathematical model, parametrized with dispersal and growth data of the of the 20 founder strains in the experiment. We find that short-term evolution was driven by selection for increased dispersal at the front and general selection for higher growth rates in all treatments. There was a good quantitative match of predicted and observed trait changes. Phenotypic divergence was mirrored by a complete genotypic divergence, indicating the highly repeatable fixation of strains that also were the most likely winners in our model. Long-term evolution in the experimental range front lines resulted in the emergence of a dispersal syndrome, namely a competition - colonisation trade-off. Altogether, both model and experiment highlight the importance of dispersal evolution as a driver of range expansions. Our study suggests that evolution at range fronts may follow predictable trajectories, at least for simple scenarios, and that predicting these dynamics may be possible from knowledge of few key parameters.

## Introduction

Predicting ecological dynamics and species’ range shifts has become a major challenge for conservation and management strategies in times of global climate and environmental change (Petchey *et al*., 2015). Indeed, whether the outcomes of range expansions or biological invasions can be predicted at all remains highly debated in ecology even in simple settings, due to the intrinsic stochasticity of these phenomena (Melbourne & Hastings, 2009; Giometto *et al*., 2014). Moreover, evolutionary processes occur at the same time scale as ecological dynamics during range expansions (Perkins *et al*., 2013; Williams *et al*., 2016), potentially exacerbating the uncertainty of outcomes (Williams *et al*., 2019).

Theory shows that range expansions can involve the concurrent evolution of dispersal and other traits (Perkins *et al*., 2013; Kubisch *et al*., 2014) and lead to the emergence of dispersal syndromes (Clobert *et al*., 2012; Cote *et al*., 2017). Individuals with greater dispersal propensity are the first to reach the range front, and they will reproduce with conspecifics that have the same fast spreader characteristics (Thomas *et al*., 2001; Hughes *et al*., 2007). Consequently, high dispersal ability and correlated life-history traits evolve in the range front populations due to spatial selection and spatially assortative mating (Phillips *et al*., 2008; Shine *et al*., 2011). Since expansion speeds are mainly influenced by dispersal and reproduction (Fisher, 1937; Kolgomorov et al. 1937), the two traits can be rapidly selected and evolve simultaneously. However, if dispersal is costly (Bonte *et al*., 2012) there may be trade-offs with other traits. Higher reproduction at the range front may come at the expense of lower competitive ability (Burton *et al*., 2010), recalling the competition-colonisation trade-off in classic species coexistence models (Calcagno *et al*., 2006).

Fast evolution in range front populations can produce eco-evolutionary feedbacks and thereby speed up the expansion process (Shine *et al*., 2011; Chuang & Peterson, 2016; Ochocki *et al*., 2019; Williams *et al*., 2019; Miller *et al*., 2020). In the emblematic example of the cane toad (*Rhinella marina*) expansion in Australia, increased dispersal at the range front coincided with evolutionary change in behavioural, morphological and demographic traits, promoting the speed of the toad expansion (Phillips *et al*., 2006; Perkins *et al*., 2013; Brown *et al*., 2014). Growing empirical evidence from other natural populations and biological systems (Simmons & Thomas, 2004; Alford *et al*., 2009; Leotard *et al*., 2009; Lombaert *et al*., 2014) suggest that dispersal evolution at range fronts is a common phenomenon. Recently, experimental evolution and microcosm landscapes have been used to test fundamental predictions (Friedenberg, 2003) and mimic range expansions in the laboratory. Experiments with ciliates (Fronhofer and Altermatt, 2015), arthropods (Ochocki & Miller, 2017; Szűcs *et al*., 2017; Weiss-Lehman *et al*., 2017; Petegem *et al*., 2018) or plants (Williams *et al*., 2016) all showed the rapid evolution of dispersal and other dispersal-related traits during the experimental range expansions. However, whether we can accurately predict these eco-evolutionary dynamics from prior information on the genetic or phenotypic characteristics of the expanding populations remains an open question.

Coupling microcosm experiments with mathematical modelling and genetic analyses provides a possible way forward to assess the predictability of range expansions (Nosil *et al*., 2020). In micro/mesocosm landscapes, we can study the repeatability of range expansions through independent replicates under controlled conditions. Using specifically tailored and parametrised mathematical models, we can then formalise putative processes of range expansion dynamics and confront predicted with observed outcomes. Genetic analysis can further characterise the degree of similarity among experimental replicates and link phenotypic trait change to genetic change.

Here, moving a step forward from previous ecological models (Melbourne & Hastings, 2009; Giometto *et al*., 2014), we employed such a combined approach to assess the predictability of evolutionary outcomes of range expansions in an aquatic model organism, the freshwater protozoan *Paramecium caudatum*. Following previous studies (Fronhofer & Altermatt (2015) Nørgaard et al. (2021), we used interconnected 2-patch systems to establish a range front treatment, where recurrent episodes of dispersal alternated with intermittent periods of population growth. In the contrasting range core treatment, only the non-dispersing individuals were maintained. We recreated these experimental treatments in a predictive mathematical model, parameterised for dispersal and growth characteristics of the 20 *Paramecium* strains that were used to assemble the founder population in the selection experiment. Based on selection from standing genetic variation in an asexual population, the model predicted the rapid divergence between range core and front populations, mainly driven by positive selection on dispersal at the front. There was a good quantitative match between model predictions and experimental results, and the most likely winner strains identified by the model corresponded to particular genotypes, found to be repeatably fixed in the experimental core and front populations. In the long run (160 dispersal/growth episodes), range core and front populations continued to diverge, resulting in the emergence of a dispersal syndrome with a competition – colonisation trade-off, hypothesised in many ecological models (Livingston *et al*., 2012). Our results suggest that, even with evolution occurring over short ecological time scales, range expansions may follow predictable trajectories and predicting these dynamics may be possible from knowledge of a few key parameters.

## Material and Methods

### Study organism and strains

*Paramecium caudatum* is a freshwater ciliate with a world-wide distribution, feeding on bacteria and detritus. Asexual reproduction occurs by mitotic division and represents the main mode of population growth. Swimming is accomplished through the coordinated movement of ciliary bands on the cell surface (Wichterman, 1986). Previous work on *P. caudatum* indicated a genetic basis of dispersal propensity, and plastic responses are induced by biotic factors, such as parasitism, chemical predator signals, or population density (Fellous *et al*., 2012; Fronhofer *et al*., 2018; Zilio *et al*., 2021). Here, we used 20 *Paramecium* strains (i.e., clonal mass cultures derived from a single individual) from various geographic origins (Weiler *et al*., 2020; Zilio *et al*., 2021) and representing different groups of mitochondrial haplotypes (“COI genotypes” or “genotypes”, hereafter; Table S1). All cultures were reared under standard laboratory conditions in lettuce medium with the food bacterium *Serratia marcescens* at 23 °C, allowing up to 3 asexual doublings per day (Nidelet & Kaltz, 2007).

### Founder strains measurements

Prior to the start of the long-term experiment, we assayed the 20 founder strains for dispersal and population growth characteristics (Table S1). For the dispersal assay, we placed aliquots of 8 mL of culture (at equilibrium density) in a 2-patch system (see below for additional details) and let the *Paramecium* disperse for 3h. Once connections were blocked, we estimated the number of residents and dispersers by taking 150-600 µL samples from the two tubes and counting the number of individuals under a dissecting microscope. Dispersal was taken as the proportion of dispersers of the total number of individuals in the system. Dispersal rates (Gaussian posteriors) were then estimated using a generalized linear mixed model (binomial error distribution; bobyqa optimizer in function glmer of R package “lme4”; Bates *et al*., 2015) for each strain with time and observation (to account for overdispersion) as random effects. We tested 4 replicates per strain. For the growth assay, we placed ca. 200 individuals (from cultures at equilibrium density) in 20 mL of fresh medium. Over the course of 6 days, we tracked population density by counting the number of individuals in daily samples of 100-200 µL. We tested 3 replicates per strain. Using a Bayesian approach (Rosenbaum *et al*., 2019), we estimated the intrinsic population growth rate (*r_0_*) and equilibrium density (*N̅*) for each replicate by fitting a Beverton-Holt population growth model to the time series data. Details of the Bayesian fitting are given in Supplementary Information (Section S1).

### Selection experiment

The selection experiment comprised a sequence of cycles, where dispersal events alternated with periods of population growth. The founder population was created by mixing the 20 strains at equal proportions in a single mass culture, which was then divided up into 15 replicate selection lines, assigned to the following three treatments. First, in the range front treatment (6 selection lines), we placed the *Paramecium* in one of the two tubes in 2-patch dispersal systems (interconnected 15-mL tubes, Fig. 1A). Connections were opened for 3 h, during which time individuals were allowed to swim to the other tube. We then collected the dispersers and cultured them for 1 week under permissive conditions in 20 mL of fresh medium (in 50-mL plastic tubes), until we initiated a new round of dispersal, again only retaining the dispersers and culturing them for 1 week, and so on. Second, the range core treatment (6 selection lines) followed the same cycles of dispersal and growth, but only the non-dispersing residents were retained after each dispersal episode. Third, in the control treatment (3 selection lines), residents and dispersers were mixed after each dispersal event and then cultured for 1 week, as in the other treatments. In corollary, the range front treatment mimics the advancing cohort of a spatially expanding population, whereas populations from the core treatment remain in place and constantly lose emigrants. The control treatment is similar to the core treatment, except for the loss of emigrants. A total of 161 cycles were accomplished. Prior to each dispersal event, ca. 1800 individuals (median; 25% / 75% quantile range: 1400 / 2700) were placed in the dispersal systems. After dispersal, the number of individuals starting the 1-week growth period were matched between treatments. Because dispersal rates were low at the beginning, these starting numbers were initially set to 200 individuals (placed in a total volume of 20 mL of fresh medium). During the following 1-week growth period, stable population sizes were typically reached within 3-4 days, with densities of ca. 240 individuals per ml (median; 25% / 75% quantiles: 180 / 360). After cycle 32, when dispersal had already reached higher levels (see Results), we adjusted the starting numbers to ca. 1500 (median; 25% / 75% quantiles: 1100 / 2000).

**Figure 1.**
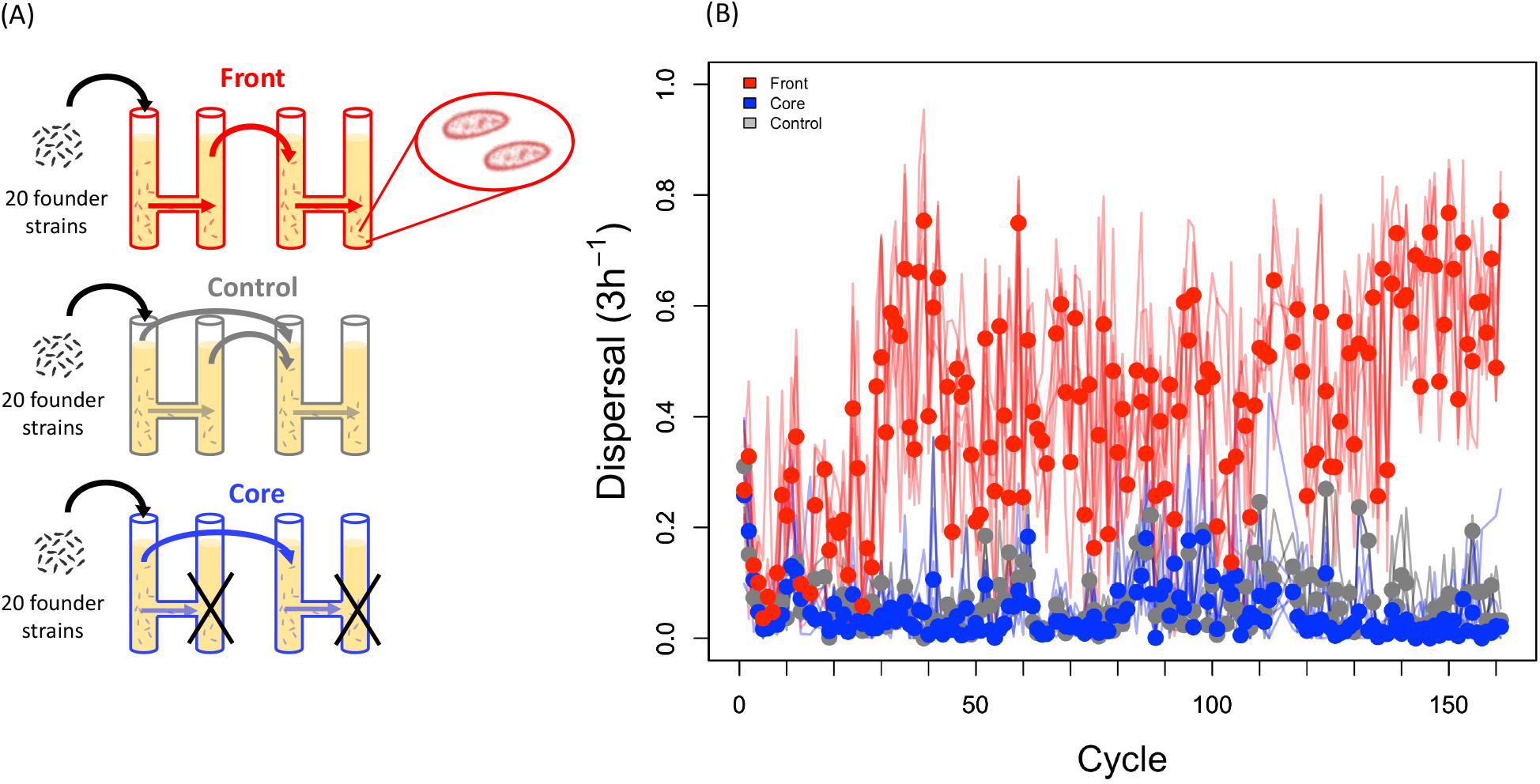
Design of range expansion selection experiment and long-term time series of dispersal in the experimental treatments. (A) Starting from a mix of 20 *Paramecium caudatum* founder strains, experimental population were allowed to disperse in 2-patch dispersal systems. For the range front treatment (red), only the dispersers were maintained and propagated for 1 week, until the next dispersal episode. In the core treatment (blue), only the non-dispersing residents were maintained at each cycle. In the control treatment, both residents and dispersers were maintained. (B) Observed levels of dispersal over the whole duration of the experiment (161 cycles, ca. 3 years). Lines show the trajectories for the individual selection lines (*n* = 15), the circles indicate the mean dispersal per treatment and cycle.

#### Data collection

For each selection line, dispersal was measured at each dispersal event and equilibrium densities (*N̅*) taken at the end of the 1-week growth period at each cycle, as described above. Furthermore, growth rate (r_0_) was determined in assays conducted at cycle 21 (year 1), 78 (year 2) and 160 (year 3), as described above, with 2-3 replicates per selection line and year. Bayesian model fitting was used to estimate *r_0_* (Section S1). Measurement of swimming behaviour were also taken in the first two years of the experiment (Section S2).

#### Genotyping

All founder strains were genotyped for the mitochondrial cytochrome *c* oxidase I (COI) gene, by extracting DNA from 10 cells per strain and applying a PCR amplification protocol and sequence analysis described in Killeen et al. (2017). At cycle 30 in the selection experiment, mixes of 50 cells from each selection line were processed in the same way and analysed for (multiple) COI marker signals. This method characterises the line for the most frequent COI genotype, and it has a resolution threshold of c. 5%, i.e., it can detect 2-3 cells of a minority genotype in the sample (Killeen *et al*., 2017). The sequences used are deposited in GenBank under the accession numbers listed in Barth et al. (2006) and Weiler *et al*. (2020).

### Range expansion model

Our model is designed to capture the specificities of the selection experiment and the characteristics of the strains in the founder population. Thus, we model the population dynamics of *Paramecium* strains, assuming logistic growth following the Verhulst equation expanded to include both intra- and inter-strain competition:

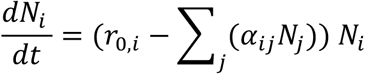

where *N_i_* is the population size of strain i, *r*_0,*i*_ is its intrinsic rate of increase and *α_ij_* as the competition coefficients. The model is parametrized with the posteriors extracted from growth curve fitting (*r_0_*, *N̅*), as described above. We make the simplifying assumption that intra- and interspecific competition is of equal strength. We further assume a quasi-extinction threshold of 0.7 (the bottleneck occurring during dispersal, we have tested the effect of different quasi-extinction thresholds ranging from 0.0001 to 0.9), which implies that strains experience an extinction if they exhibit densities below this value.

We model the community dynamics of the strains for 7 days, followed by a 3h dispersal phase in a 2-patch metapopulation. During this 3h dispersal phase all strains can disperse from their patch of origin to their destination patch according to the dispersal rates estimated from dispersal assay; the model is parametrized with posteriors extracted from the statistical analysis described above. After the dispersal phase we follow the patch of origin (residents in the range core treatment), the destination patch (dispersers in the front treatment) or the combined patches (dispersers and residents mixed in the control treatment). We repeat this procedure for a total of 10 iterations. As in the experiment, we control for densities between rounds of iteration by selecting the equivalent of 10 mL samples.

This approach allows us to predict, based only on measurements of growth parameters and dispersal rates of the founder strains, which strains should predominate in each of the three treatments at the end of the experiment. It is important to keep in mind that the underlying model is deterministic. However, since we parametrize the model with draws from posteriors, our approach takes into account the uncertainty associated with the data and yields a distribution of likely outcomes, given these uncertainties. Note that our model depicts a scenario of selection from standing genetic variation; it does not include mutational change.

### Statistical analysis

Statistical analyses were performed in R (ver. 4.1.2; R CoreTeam 2021 and JMP 14 (SAS Institute Inc. 2018). We analysed dispersal (proportion of dispersers), using generalised linear models (GLM) with binomial error distribution. We considered selection treatment (core, front, control), experimental cycle and selection line (nested within treatment) as explanatory variables. We analysed variation in intrinsic population growth rate (*r_0_*) and equilibrium density (*N̅*; averages per selection line and year) using GLMs, with selection treatment, year and selection line as explanatory variables. To illustrate how selection acts on standing genetic variation in our model, we associated the winning probability of each of the 20 founder strains with their respective median values of dispersal, r*_0_*, and *N̅* from the distributions used by the model. For each treatment, we then performed multiple regressions, with winning probability as response variable and the three traits as explanatory variables. To investigate associations between dispersal, *r_0_* and *N̅*, we constructed a data matrix based on trait means per year and selection line (3 years x 15 selection lines, *n* = 45), after centering and scaling trait distributions. One range-front line was lost in year 3, leading to *n* = 44. We used a Bayesian approach with the “rstan” package version 2.21.2 (Carpenter *et al*., 2017) to estimate pairwise trait correlations (see section S3). We also performed a principal component analysis (PCA), considering the joint variation in all three traits.

## Results

### Predicted and observed short-term trait evolution

Over the first 25 cycles of the selection experiment, we observed a strong increase in dispersal in the range front treatment. Dispersal reached 22.3% (± 0.012 SE, averaged over cycles 15-25) at the front, compared to only 4.4% (± 0.004 SE) in the core treatment and 6 % (± 0.012 SE) in the control treatment (effect of selection treatment: χ_2_^2^ = 119.7; p < 0.001). Increased dispersal at the front established within only a few cycles, and was formally significant for the first time at cycle 8 (cycle-by-cycle analysis: p < 0.001). Our parametrized model captured this rapid increase of dispersal in the range front treatment (Fig. 1B), and there was a quantitative match between the distribution of endpoint levels of dispersal in the model and observed values in the experiments (Fig. 2A). The model further predicted general increases in growth rate (r*_0_*) and equilibrium density (*N̅*) in all treatments, from 0.07 in the ancestral mix to 0.08 in core and front end-point populations. This is consistent with results from the growth assay conducted at cycle 21, where estimates of r*_0_* for the 15 selection lines are well within the central range of predicted values in the model (Fig. 2B). As predicted, selection treatments did not significantly differ in r*_0_* (treatment: F_2,12_ = 1.2; p = 0.354). Unlike in the model, range front lines produced nearly 20% higher equilibrium densities than did range core and control lines (treatment: F_2,12_ = 11.1; p = 0.003).

**Figure 2.**
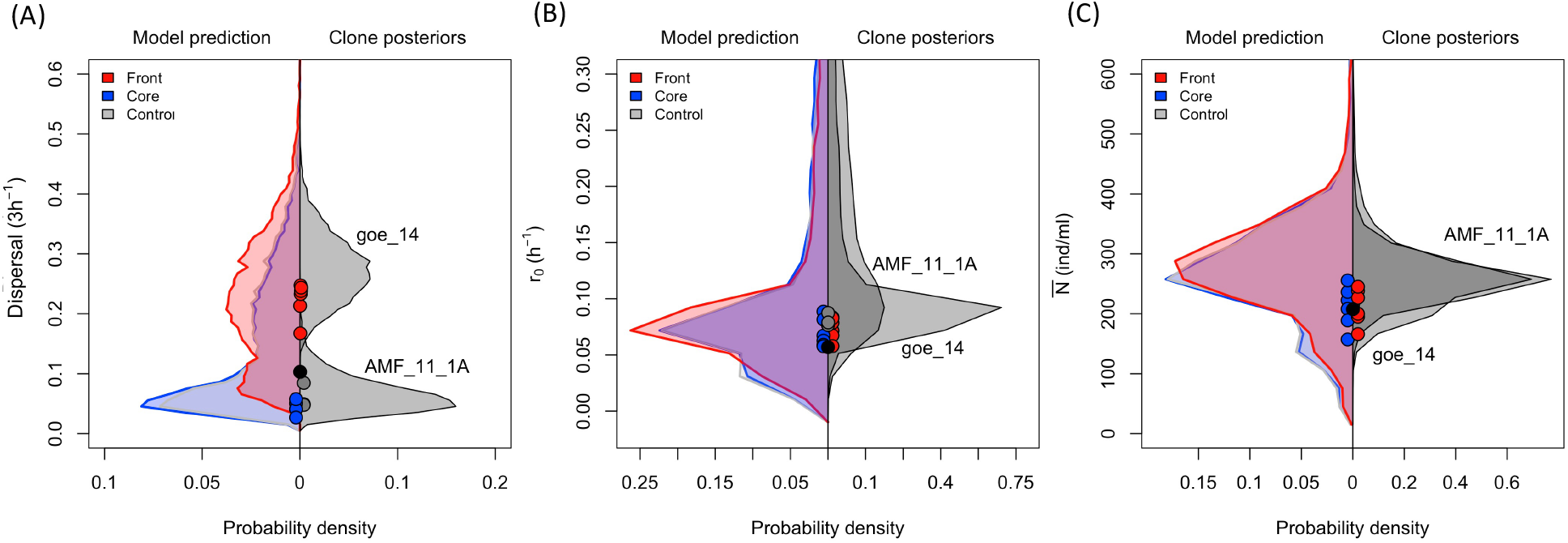
Model endpoint predictions for (A) dispersal, (B) growth rate (*r0*), and (C) equilibrium density (*N̅*). In each panel, left: model predictions for the 3 treatments; right: posteriors distributions of the most likely winner strains in the range core (AMF_11_11A) and range front (goe_14) treatment. Circles are the average values measured for each experimental selection lines after 15-25 cycles (“short term”). Different colours represent the different treatments. The black circles represent the ancestral means (founder population).

### Predicted and observed short-term changes in strain composition

Our model finds strong variation in the fixation probability among the 20 strains, and different treatments have different most likely winners (range: 0.7-16.8 %; Fig. S7).

For the range front treatment, multiple regression analysis (Table S1) shows that both dispersal and r*_0_* are positively associated with strain winning probability, and this with equal strength (standardised beta (β) regression coefficients: +0.55 and +0.64, respectively; Fig. 3). Thus, selection is predicted to favour strains that both disperse more and grow faster. In contrast, in the core and control treatments, strain winning probability is mainly associated with high growth rate (β > +0.96), accompanied by weak selection against clones with higher dispersal (β ≤ -0.27) or equilibrium density (β ≤ -0.33).

**Figure 3.**
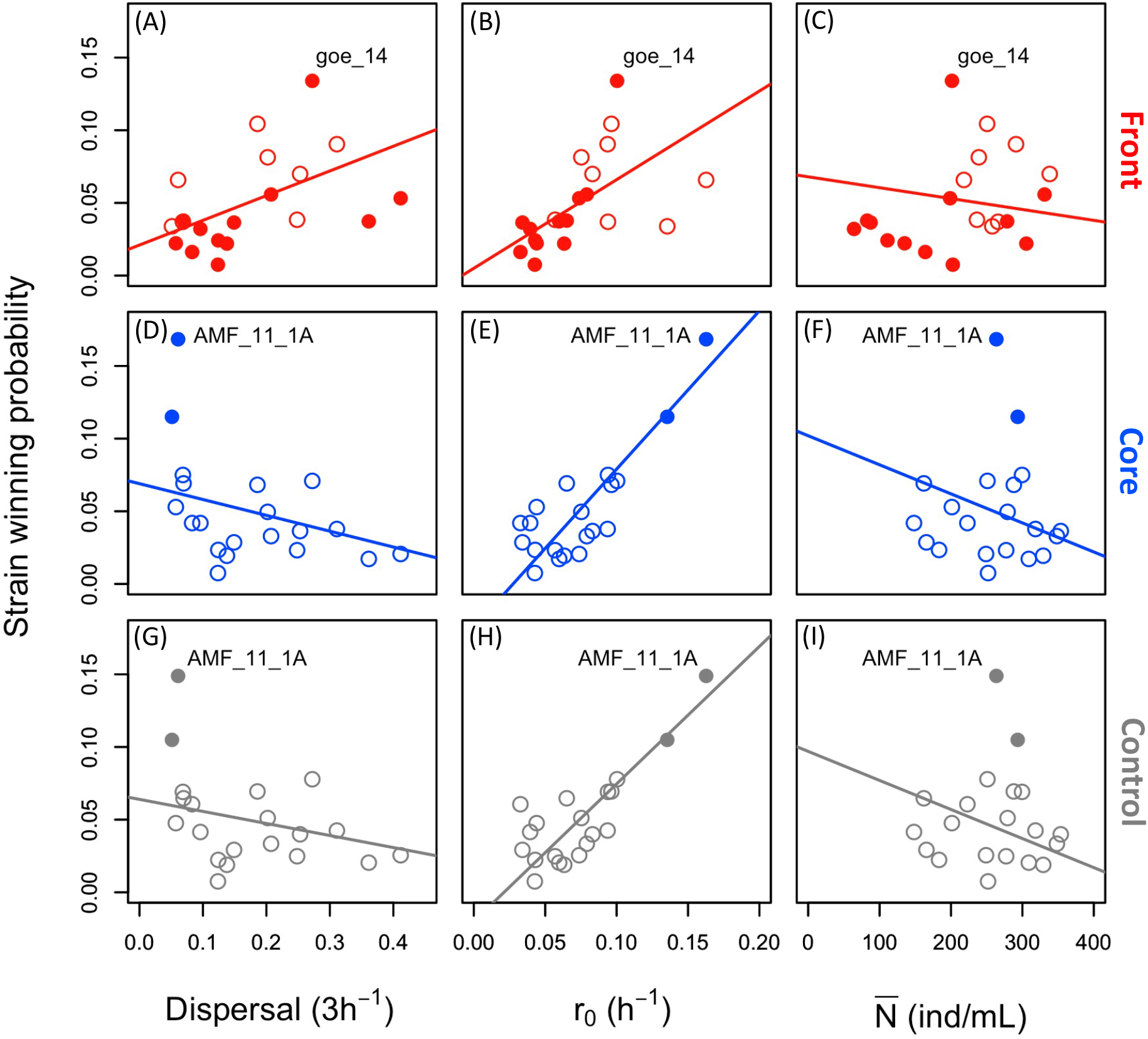
Winning probability (frequency of going to fixation in 10k model runs) of each of 20 strains from the founder population, as a function of its dispersal, growth rate (*r0*) and equilibrium density (*N̅*), shown for range front (A-C), range core (D-F) and control (G-I) treatments. Full circles denote the potentially fixed and open circles the eliminated strains, according to genetic analysis (COI genotype). Regression lines obtained from multiple regression models. Different colours represent the different treatments.

Molecular analysis of the 15 selection lines indicates complete genetic divergence between selection treatments. For all 9 range core and control lines, only the b05 COI genotype was detected. The two strains in the founder population that carry this genotype (Table S1) have very high growth rates and very low dispersal, the trait combination favoured in the model. Indeed, the candidate strain AMF_11_1A has the highest growth rate overall and is the most likely winner in core and control treatments according to our model (Fig. 3). In contrast, all 6 range front selection lines appear to be fixed for the b07 COI genotype. This genotype is shared by 13 founder strains (Table S1), which may thus have gone to fixation in groups or individually. Among these candidate strains is the most likely winner (goe_14) predicted by the model: it has the highest growth rate and the third-highest dispersal, in line with the prediction of the two traits being under joint positive selection in this treatment. As shown in Fig. 2, trait values of the most likely front and core winner strains (goe14 vs AMF_11_1A; strain posterior distributions on the right) show a good match with both the predicted model outcomes (distributions on the left; Fig. 2) and the experimental data.

### Long-term changes

#### Dispersal

In addition to the short-term evolution (see above), we also observed a long-term increase in dispersal in the range front treatment over the entire time span of the three years of the experiment (cycle x treatment interaction: χ_2_^2^ = 88.8; p < 0.001; Fig. 1). This trend is significant, even when omitting the first 50 cycles (χ_2_^2^ = 51.7; p < 0.001). We found little evidence for a dispersal difference between range core and control lines, neither overall (contrast core vs control: p > 0.68) nor when considering individual cycles (11 cycle-by-cycle contrasts with 0.0078 < p < 0.09, none significant after correction for multiple testing).

#### Demographic traits

While no significant treatment effects were detected in the first growth assay (cycle 21, see above), range front lines had nearly 2-fold lower values of r*_0_* than range core lines in assays conducted in year 2 and 3 (year x treatment: F_4_ = 6.66; p < 0.001; Fig. SI 1A). Furthermore, while beginning to grow more slowly, range front lines continued to produce up to 2-fold higher *N̅* than range core and control lines (treatment: F_2_ = 34.21; p < 0.001; Fig. SI 1B).

#### Trait associations

Fig. 4A-C illustrates short-and long-term trends in pairwise trait associations, in relation to the model predictions. For dispersal and r*_0_* (Fig. 4A), there was no clear relationship between the two traits after short-term selection (year 1). However, in year 2 and 3, observed data points tend to fall outside the main predicted ranges, and a negative relationship between dispersal and r*_0_* emerged (Fig. 4A). This negative association is highly significant over all lines and years combined (r = -0.627, 95% CI [-0.771; -0.434]), but also holds for year 2 and 3 separately (Fig. S3). The positive relationship between dispersal and *N̅*, already observed as a short-term trend, further consolidated in year 2 and 3 (Fig. 4B), again with values mostly falling outside the main predicted short-term ranges. The correlation is significant overall (r = 0.599, 95% CI [0.347; 0.725]), as well as for each year separately (Fig. S3). Furthermore, diverging trends in core and front selection lines lead to a negative association between r*_0_* and *N̅* (Fig 4C). The negative correlation is of intermediate effect size overall (r = -0.325, 95% CI [-0.575; -0.031]), and is significant in all three years separately (Fig. S3). It should be noted that all of these main trends of divergence hold, when we correct for year effects, by expressing front and core line data relative to the control treatment in each year (Fig. S5).

**Figure 4.**
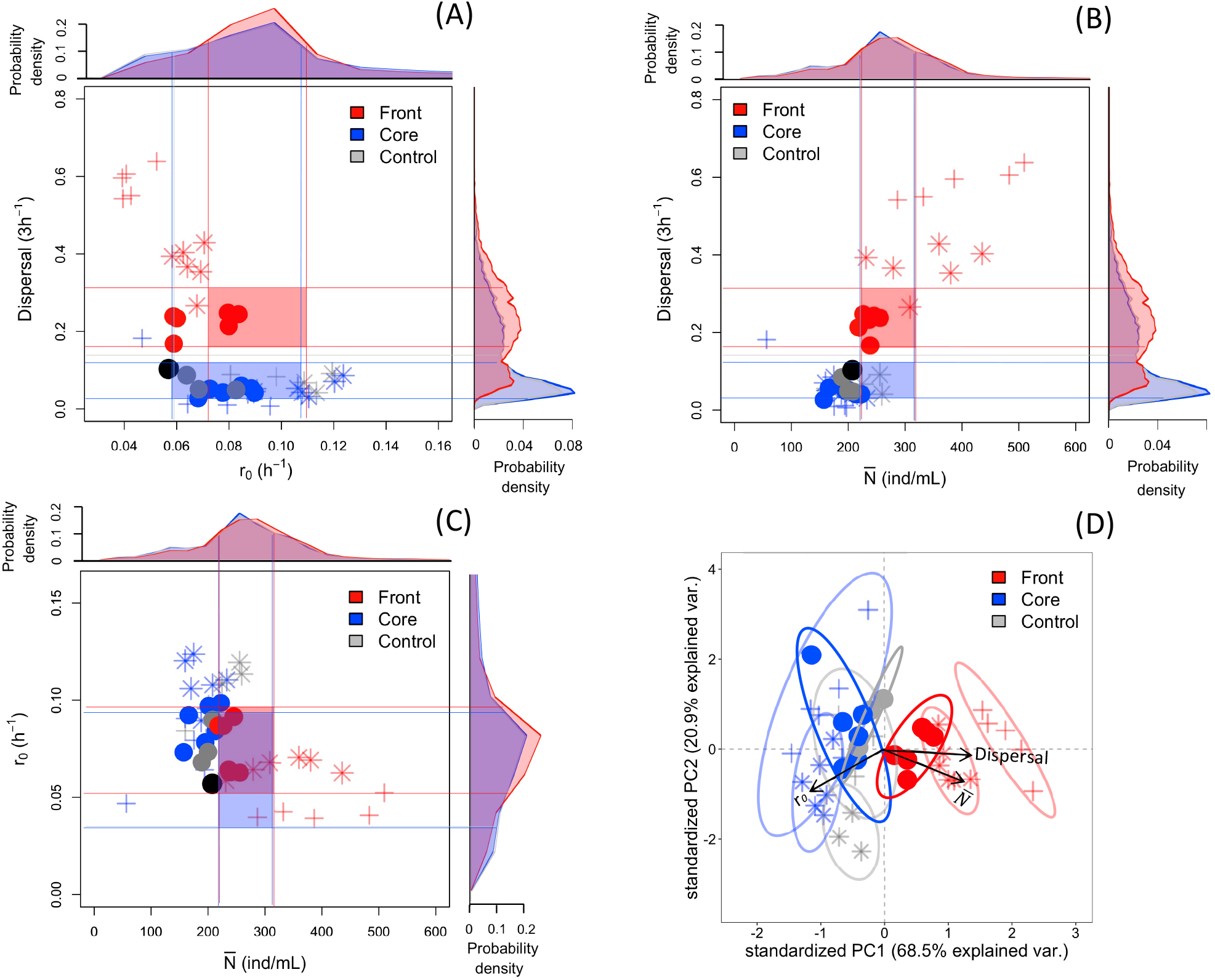
Short- and long-term traits associations observed in the experiment, in relation to short-term predictions in the model. (A-C) Bivariate correlations between dispersal, growth rate (*r0*) and equilibrium density (*N̅*). Circles are the average values measured for each experimental selection line in year 1 (cycles 15-25; “short term”). Stars refer to year 2 (cycles 74-84) and cross symbols to year 3 (cycles 154-161). From the distributions of the model predictions (outer part of graphs), the central range of each trait (50% high density probability interval, HDPI; thin lines) can be defined; the overlap zones of the HDPI (shaded square areas) represent the predicted trait association for each treatment, after short-term evolution. Observations falling outside of the overlap areas indicate deviation from the model, possibly due to de novo evolution (year 2 and 3). The black circles represent the mean ancestral trait association (founder population). (D) Principal Component Analysis (PCA) of all three traits combined according to the first two principal components of the PCA. The arrow length represents the loading value of the trait, while opposite arrow direction indicates opposite trend between traits. Different symbols correspond to the different years (circles, year 1; stars, year 2; cross, year 3). The ellipses are the 95 % containment probability region per treatments and year. Different colours represent the different treatments.

Principal Component Analysis (PCA, Fig. 4D) summarises the patterns of phenotypic divergence. Demography-related traits and dispersal are pulling in approximately equal strength on PC axis 1, but in opposite directions (PC 1 loadings: *r_0_* = -0.53; *N̅* = +0.57; dispersal = +0.62). Thus, range front lines are characterised by a combination of higher equilibrium density and dispersal, but lower intrinsic population growth rate relative to range-core and control lines (MANOVA: F_2,37_ = 10.85, p < 0.001). The separation of clouds indicates the progressive divergence through time, with a maximum in year 3. There is little differentiation between range core and control treatments.

## Discussion

Predicting range expansions with ecology and evolution occurring on the same timescale is a challenging task. Building on previous ecological range expansion studies (Melbourne & Hastings, 2009; Giometto *et al*., 2014), we included short-term evolution in a simple model parameterised for our laboratory system and confronted predicted evolutionary outcomes with results from experimental range expansions. Both model and experiment show rapid divergence between range core and front treatments, with selection for higher dispersal at the front. The repeated fixation of particular COI genotypes in the experimental lines corresponded to strains identified as most likely winners in the model. This match between predicted and observed outcomes suggest a certain predictability of range expansions, even when evolutionary change occurs. Over longer time scales, experimental range core and front populations continued to diverge, indicating *de novo* evolution and resulting in the emergence of dispersal syndromes.

### Dispersal and growth rate are main targets of selection

In the context of reaction-diffusion models, dispersal (diffusion) and population growth at low densities are the two key traits for understanding and predicting range expansion dynamics (Fisher, 1937; Kolgomorov et al. 1937). Consistent with this view and previous studies (Phillips *et al*., 2010; Shine *et al*., 2011), dispersal and population growth were here identified as main targets of selection.

Higher dispersal was immediately selected from standing genetic variation at the range front and weakly selected against in the range core in the model as well as in the experiment, where range front populations showed increased dispersal already after the first few cycles. Such strong and fast selection on dispersal in the vanguard front populations has been found in similar experiments (Fronhofer & Altermatt, 2015; Williams *et al*., 2016; Ochocki & Miller, 2017; Szűcs *et al*., 2017; Weiss-Lehman *et al*., 2017; Petegem *et al*., 2018), but also in natural populations (Phillips at al. 2006; Perkins et al. 2013). Dispersal evolution might therefore accelerate the speed of range expansion already over very short time scales (Ochoki & Miller, 2017; Miller et al. 2020).

Contrary to more standard views of range expansion with r- and K-selection (Charlesworth, 1971; Roughgarden, 1971; Burton *et al*., 2010), growth rate was under positive short-term selection in both range core and front treatments. This can be explained by the fact that populations in all treatments experienced regular bottlenecks, thus imposing general selection for increased growth rate, a trait for which there was ample variation among founder strains (Fig. S4 A-C). Importantly, however, our model shows that dispersal and growth rate can be simultaneously selected in the range front treatment (Fig. 3). Whether one or the other trait has more weight depends on the stochasticity introduced by the dispersal bottlenecks, implemented in the model via the quasi-extinction threshold. With small bottlenecks, even weak dispersers make it into the new patch and can subsequently regrow to high density. Indeed, additional model scenarios show that when we decrease the quasi-extinction threshold, selection for growth rate overrides selection for dispersal and the strain with the highest growth rate becomes fixed in all treatments (Section S8). However, the model scenario that fits the observed data indicates a large enough extinction threshold in our experiment, putting equal selective weight on dispersal and growth rate (Fig. 2) and allowing selection to pick the best possible disperser strain that still has a high growth rate.

### Predictability of outcomes

Genetic analysis indicates that the experimental selection lines became fixed for single COI genotypes. Despite limited resolution (several strains have the same COI), there is a good correspondence with model predictions: Range core and control treatments were fixed for the b05 COI genotype, and the two b05 strains in the founder population were the most likely winners in the model, due to their particularly high growth rate (Table S1). In the range front treatment, there is more uncertainty (12 strains carry the b07 genotype fixed in this treatment), but among the possible candidates only the predicted most likely winning strain (goe_14) has both high dispersal and high growth rate (Fig. 3). Additional sequencing would be required to determine whether these selection lines are fixed for the same or different (combinations of) strains.

Although our model seems to correctly identify the most likely winner strains, it nonetheless predicts the frequent fixation of strains with alternative COI genotypes (Fig. 3). Indeed, according to the model, our exclusive finding of b05 strains in all 9 range core and control lines is highly unlikely (0.28^9^ < 0.0001). Similarly, even in the front treatment, the expected probability of exclusive fixation of b07 strain(s) is well below 5% (0.48^6^ = 0.01). In this sense, our experiment was more deterministic than the model. Possibly, when we determined dispersal and growth of the individual strains, a large measurement error was added to biologically relevant variation (Fig. S4). This additional noise then cascades through the model, from the strain posterior distributions (making them wider) to the phenotypic composition of the founder population (making strains more similar) to model outcomes (making them more variable). Alternatively, our model may be missing additional factors, such as direct strain-strain interactions or density-dependent dispersal, which potentially amplify among-strain variation in performance. Regardless, one main conclusion from this model-experiment confrontation is that evolution can be fairly predictable, at least in the short term. As already shown for ecological models (Giometto *et al*., 2014), realistic predictions can indeed be made about range expansion dynamics, at least in highly controlled laboratory settings. Here we infer trait change from knowledge of standing genetic variation in only a few parameters, suggesting that such models can be readily extended to include evolution.

### Long term evolution of dispersal syndromes and emergence of trade-offs

Experimental evolution studies show that adaptation to novel conditions may reduce performance in other environments (Kassen, 2014). The emergence of such trade-offs depends on underlying biochemical and life-history constraints (Walsh & Blows, 2009), but also on historical contingency, determining the composition and genetic architecture of the ancestral population, and thus the available trait space for selection to act on.

In our case, short-term selection from standing genetic variation did not seem to produce clear trade-offs. In the long run, however, range front and core populations continued to diverge in multiple traits (Fig. 4D), and the increase in dispersal in the front treatment was associated with a decrease in growth rate (Fig. 4A). Such coupled responses in dispersal and life-history traits are referred to as dispersal syndrome (Clobert *et al*., 2012; Cote *et al*., 2017). Typically, they involve the emergence of a competition-colonisation trade-off, where dispersal evolution coincides with selection for opportunistic growth strategies (r-selection). Theoretical and empirical studies have demonstrated the importance of dispersal syndromes in generating eco-evolutionary feedbacks and accelerating the pace of range expansions and biological invasions (Burton *et al*., 2010; Perkins *et al*., 2013; Ochocki *et al*., 2019; Miller *et al*., 2020).

Dispersal - growth trade-offs were previously reported for this (Zilio *et al*., 2020) and another ciliate species (Fronhofer & Altermatt 2015). In these systems, growth rate is a good indicator of competitive ability, and the trade-off with dispersal likely reflects a true life-history constraint, mediated through energy costs of foraging activity (Fronhofer & Altermatt 2015).

The evolved differences between core and front lines are stable, even after switching core and front treatments for multiple cycles (Fig. S6.1). Moreover, mixes of core and front lines readily respond to dispersal selection (Fig. S6.2), making new selection experiments possible, where phenotypic measurements change can easily be combined with the tracking of COI genotype frequencies.

### Advantages and limitations of an asexual reproduction scenario

In this study, we consider asexual reproduction in both model and experiment. Hence advantageous allele combinations are not broken apart or reshuffled by sex and recombination (Otto, 2009; Lehtonen *et al*., 2012), such that strains with favourable trait combinations rapidly increase in frequency in our range and core treatments. Similar results were reported for experimental range expansions of the plant *Arabidopsis thaliana*, where the fastest-dispersing clonal genotype became predominant in multiple replicate selection lines, all starting from the same initial mix of clones (Williams *et al*., 2016). Thus, asexual reproduction narrows down the variability in the range expansion outcomes and, as we show here, makes predictions possible with relatively simple models.

Clearly, recombination will make predictions more difficult, and replicated range expansion experiments with sexually reproducing organisms already showed higher variability and uncertainty in final outcomes (Ochocki & Miller, 2017; Weiss-Lehman *et al*., 2017; Petegem *et al*., 2018). For example, recombination may slow down range expansions in the short term, but speed up longer-term responses by creating novel trait associations not previously available. In our system, sex may have immediate and strong fitness consequences due to the nuclear dimorphism typical of all ciliates. Aside from creating novel genetic variants (in the germline micronucleus), sexual reproduction also involves the recreation of a new somatic macronucleus and thereby the loss of any (somatic) adaptation acquired during asexual life (Verdonck *et al*., 2021).

### Conclusions

Predicting evolution is arduous because of the intrinsic tension between determinism and contingency (Blount *et al*., 2018), and it demands an adequate theoretical representation of the eco-evolutionary processes in the biological system in question and reliable information on the genetic variation in the relevant traits (Nosil *et al*., 2020), as we describe in this work. At least in simple settings as ours, accurate predictions of the evolutionary outcomes of range expansions require surprisingly few parameters, and independent biological realisations can be highly repeatable. Future studies will need to consider, for example, more realistic landscape scenarios and interactions with other species occurring during range expansion. This would imply a more systems-biology approach, with simulations calibrated on the empirical knowledge of the specific ecological scenario and biological players (Papp *et al*., 2011). More generally, increasing our capacities to make reliable quantitative predictions of invasive eco-evolutionary processes is critical to a variety of issues, from conservation and biocontrol strategies to antibiotic development and disease management.

## Acknowledgements

GZ was supported by a grant from the Agence Nationale de Recherche (n° ANR-20-CE02-0023-01) to OK. This is publication ISEM-XXXX-XXX of the Institut des Sciences de l’Evolution. Florent Deshors and Thomas Teissedre assisted with movement and growth assays. Alison Duncan, Flore Zelé and Alexey Potekhin gave helpful comments for the experimental set up and the interpretation of the results.

## Author Contributions section

OK conceived the study. OK and CGB performed the experimental work. SK performed the molecular analyses. EAF built the model. GZ, EAF and OK performed the statistical analysis and interpreted the results. GZ, EAF and OK wrote the first draft of the manuscript and all authors commented on the final version.

## Data availability statement

The data and model will be made available upon potential acceptance (via Dryad/Figshare repository).

## Supplementary Information

### S1 Growth characteristics of long-term selection lines: *r_0_* and *N̅*

In growth assays, intrinsic population growth rate (*r_0_)* was measured after 21 (year 1), 78 (year 2) and 160 (year 3) dispersal / growth cycles. Equilibrium density (*N̅*) was taken for each selection line at the end of the 1-week growth period at each cycle during the long-term experiment. Here we averaged *N̅* for each line over 9-10 cycles in year 1 (cycle 15-25), year 2 (74-84) and year 3 (154-163).

**Figure S1.**
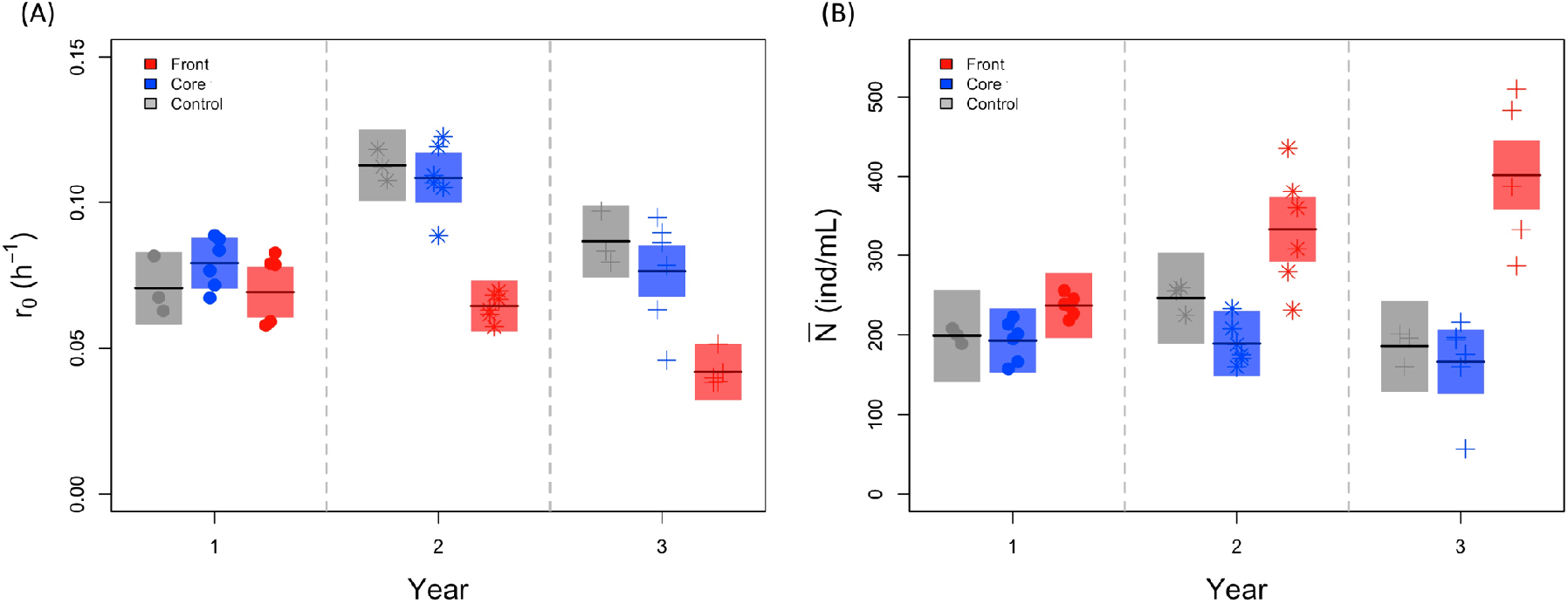
(A) Intrinsic population growth rate (*r0*) and (B) mean density at the end of the cycle ( ) from core, front and control treatments (respectively in blue, red and grey). Full symbols represent the mean values for each selection line (*n* = 15), different symbols (circle, star, cross) refer to the three different years where measurement were taken. Shaded panels show means and 95 % confidence intervals of the model predictions.

### Script Beverton-Holt fitting (*r_0_*)

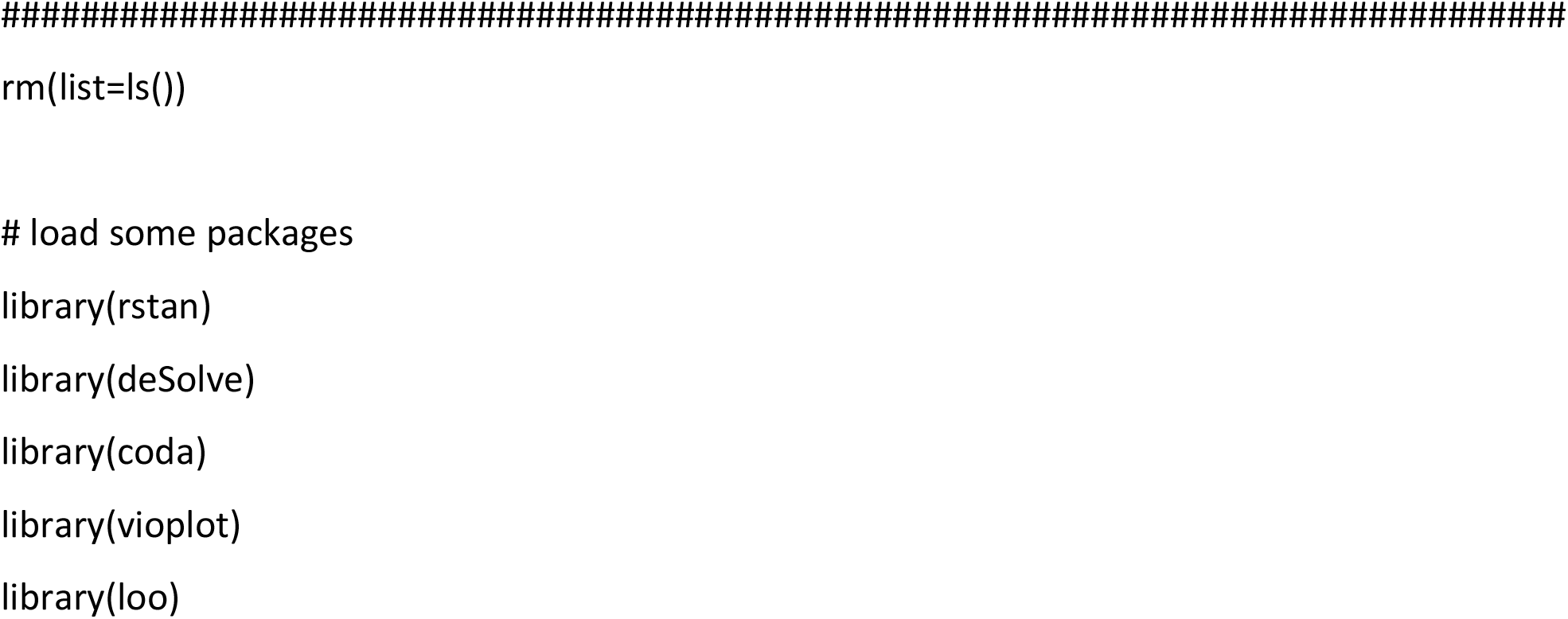
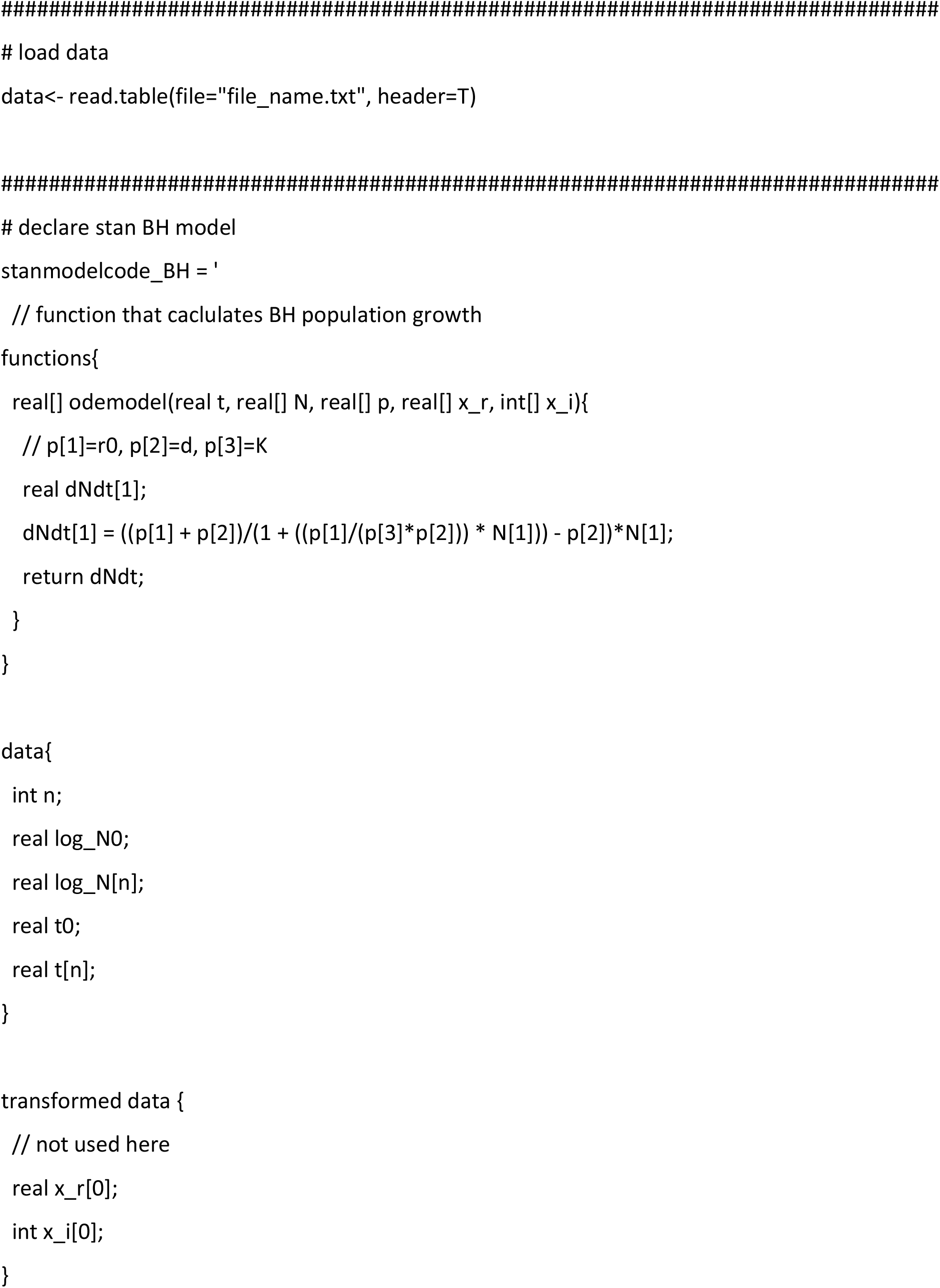
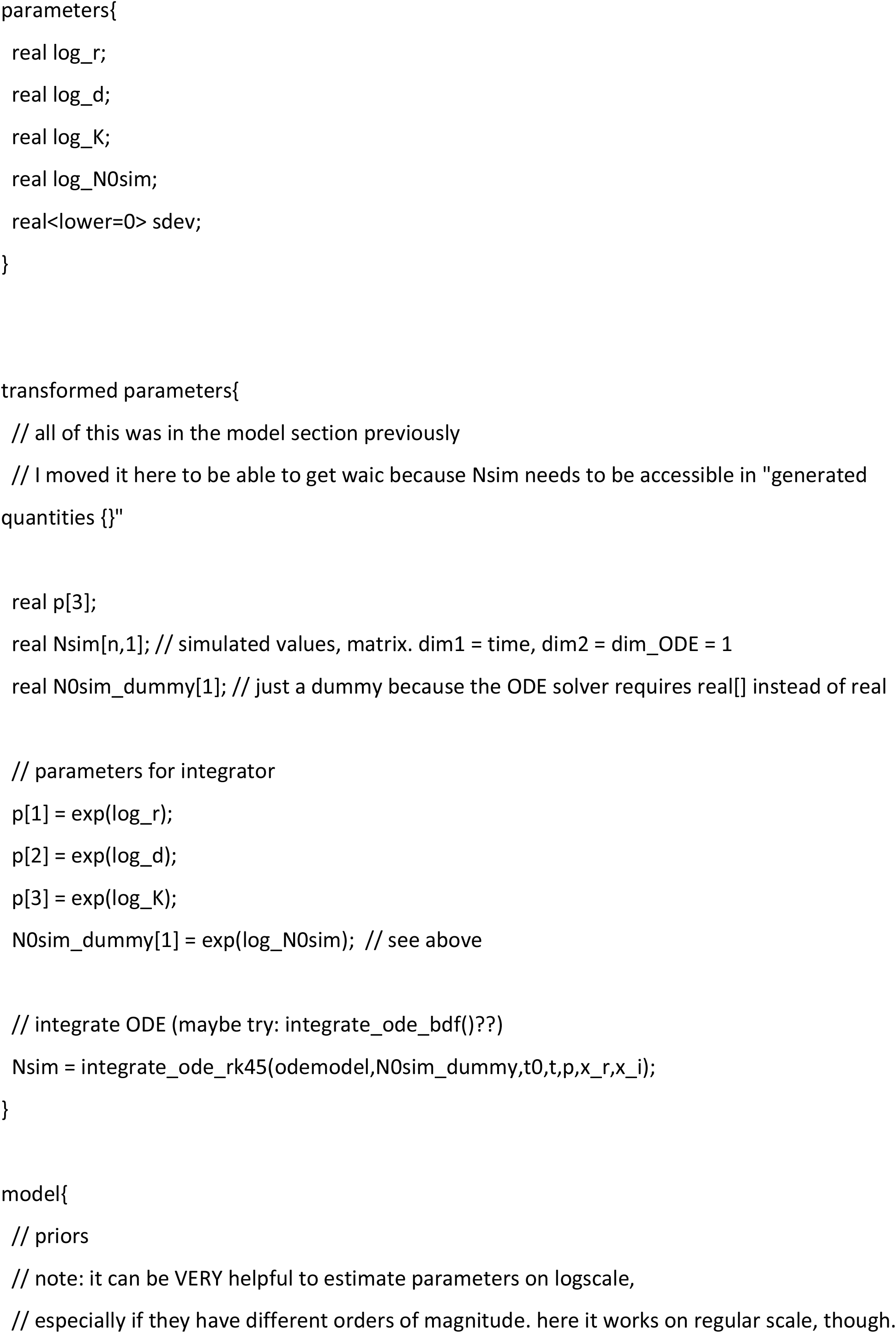
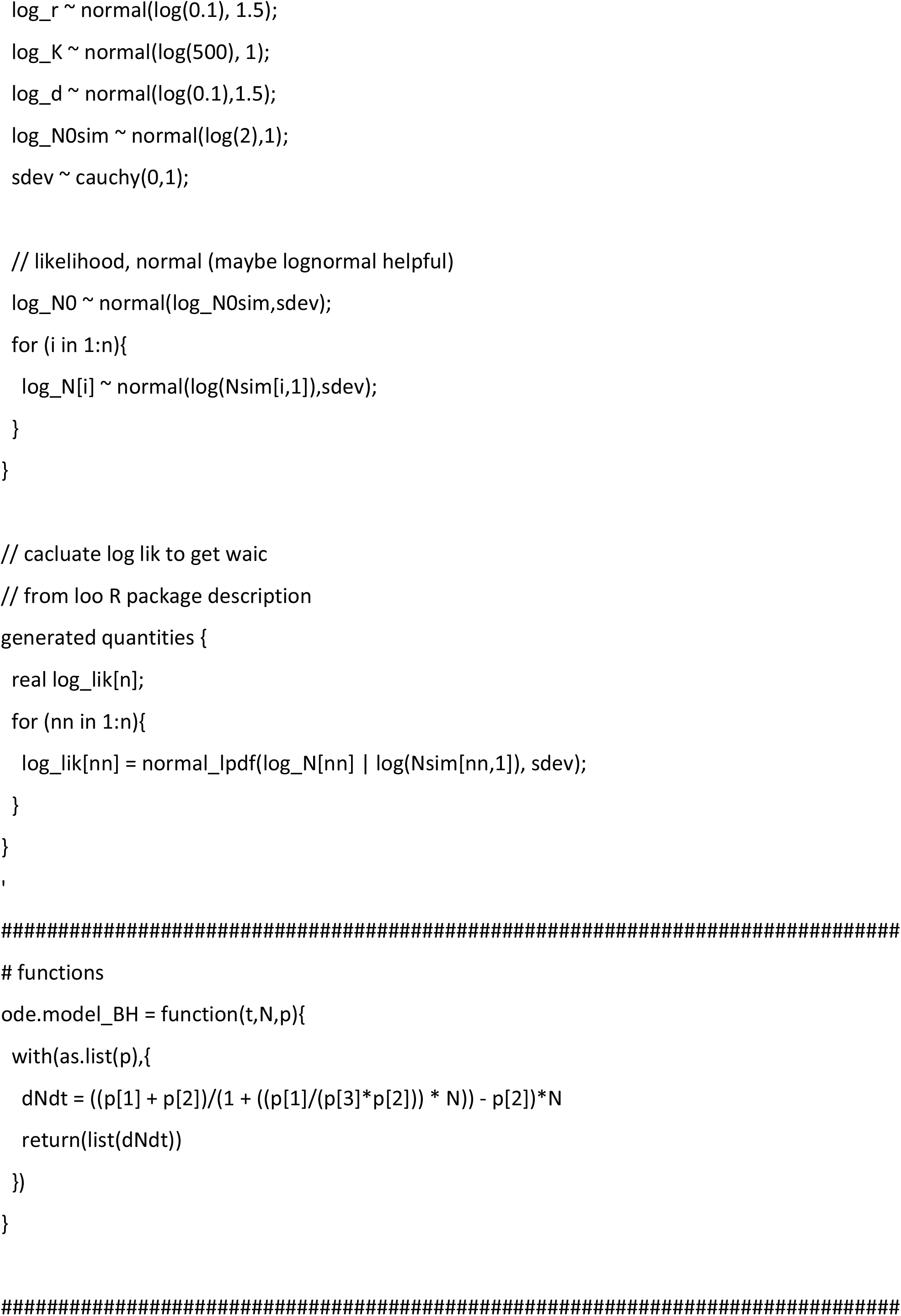
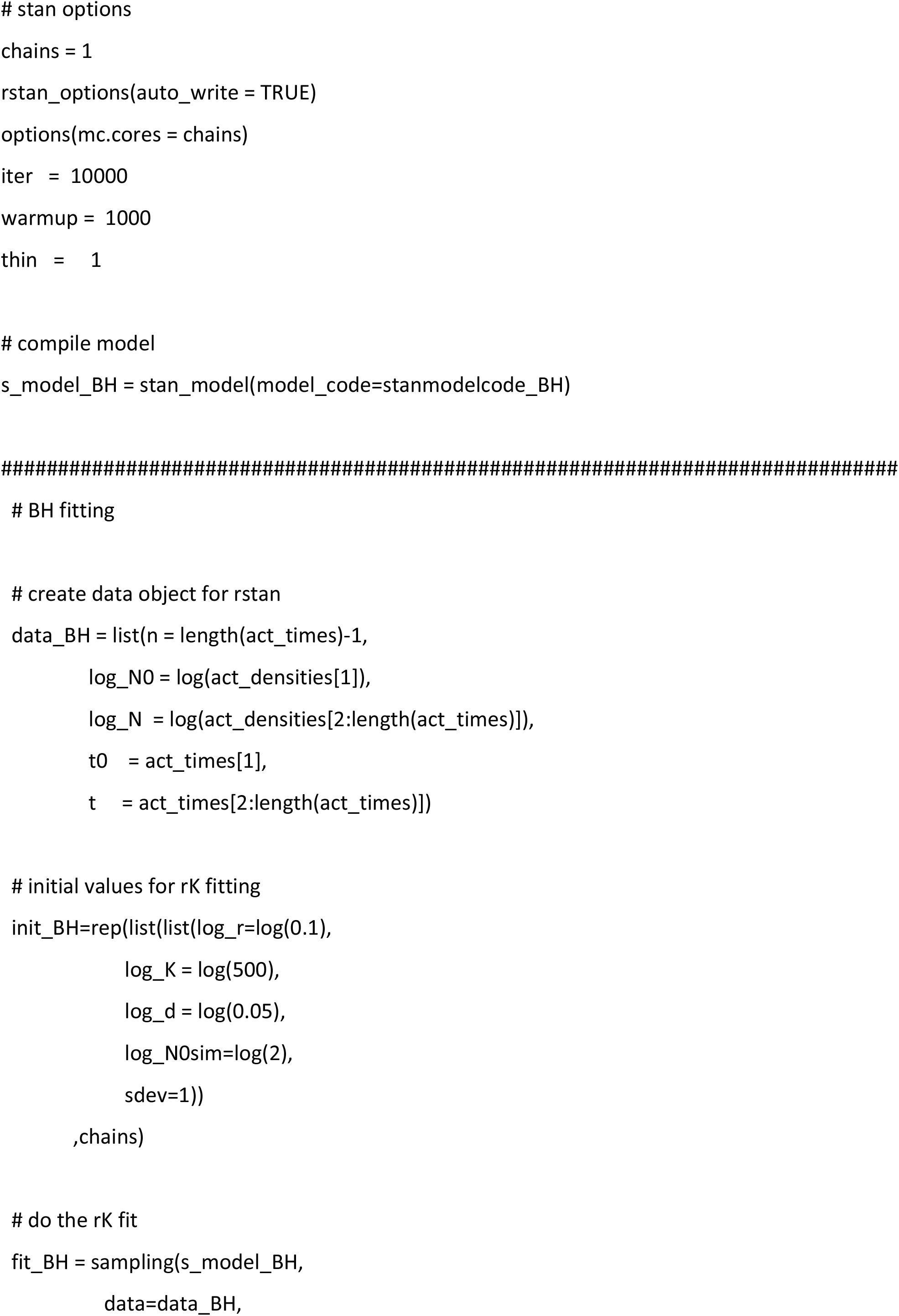
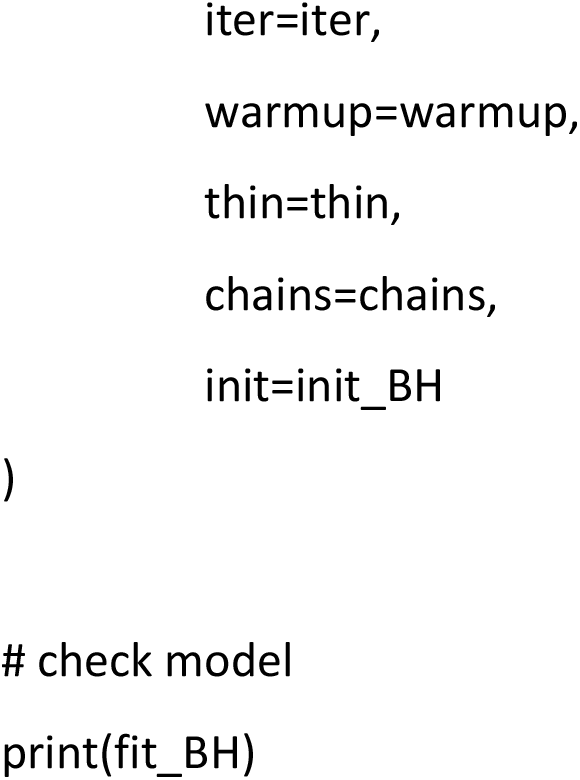

### S2 Swimming behaviour

We collected data on swimming behaviour for each selection line in year 1 (cycles 53, 54, 55, 58, 74) and year 2 (cycles 103 and 116), using an automated video analysis pipeline (Fronhofer & Altermatt, 2015; Pennekamp *et al*., 2015). To this end, 120-µL samples (ca. 20 individuals) from populations at equilibrium were placed on a microscope slide and videos were recorded under a stereomicroscope (Perfex SC38800 camera; settings: frames per second: 15; duration: 5 s; total magnification: 10x). One video per selection line and cycle was recorded, except for cycle 74 (n=4). We analysed the videos using the “bemovi” package (Pennekamp *et al*., 2015, see script below), which provided estimates of individual *Paramecium* swimming speed and the tortuosity of swimming trajectories, an indicator of changes in the swimming direction (standard deviation of the turning angle distribution). For analysis, averages of swimming speed and tortuosity were calculated for each sample.

#### Results

*Paramecium* from range front lines had a significantly lower swimming speed (-41%) than those from the range core and control treatments (treatment: F_2,12_ = 33.5; p < 0.001; Fig. S2). Across selection lines and years combined, swimming speed in the assays was negatively correlated with dispersal rate observed in the selection lines (r = -0.72, n = 30, p < 0.0001), meaning that lines with a higher dispersal generally had a lower swimming speed. Tortuosity of swimming trajectories were negatively correlated with swimming speed (all replicates: r = -0.29, n=135, p = 0.0005), but did not significantly differ among treatments (p > 0.545).

**Figure S2.**
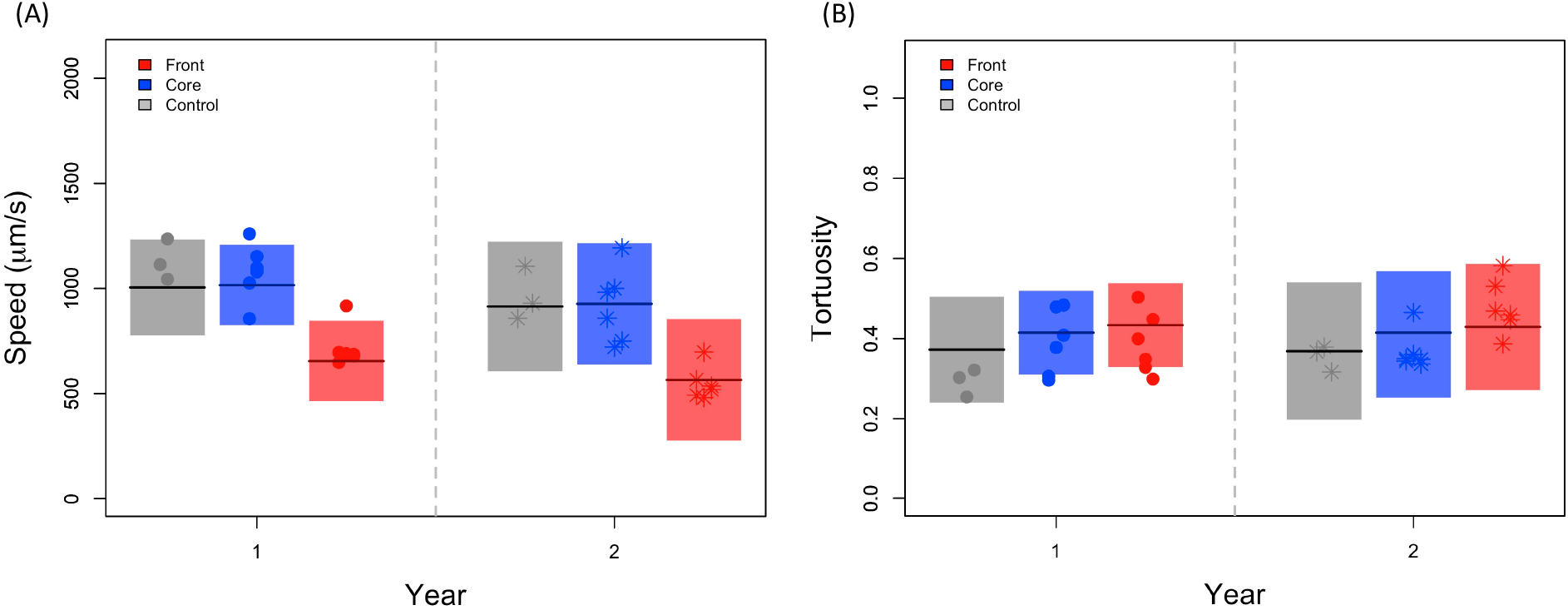
Estimates of the (A) swimming speed and (B) tortuosity of swimming trajectories from core (blue), front (red) and control (grey) treatments. Full symbols represent the mean values for each selection line (*n* = 15), with different symbols (point, star) corresponding to the first two years of the study. No measurements were taken in the year 3. Shaded panels show means and 95 % confidence intervals of the model predictions.

### Script video analysis

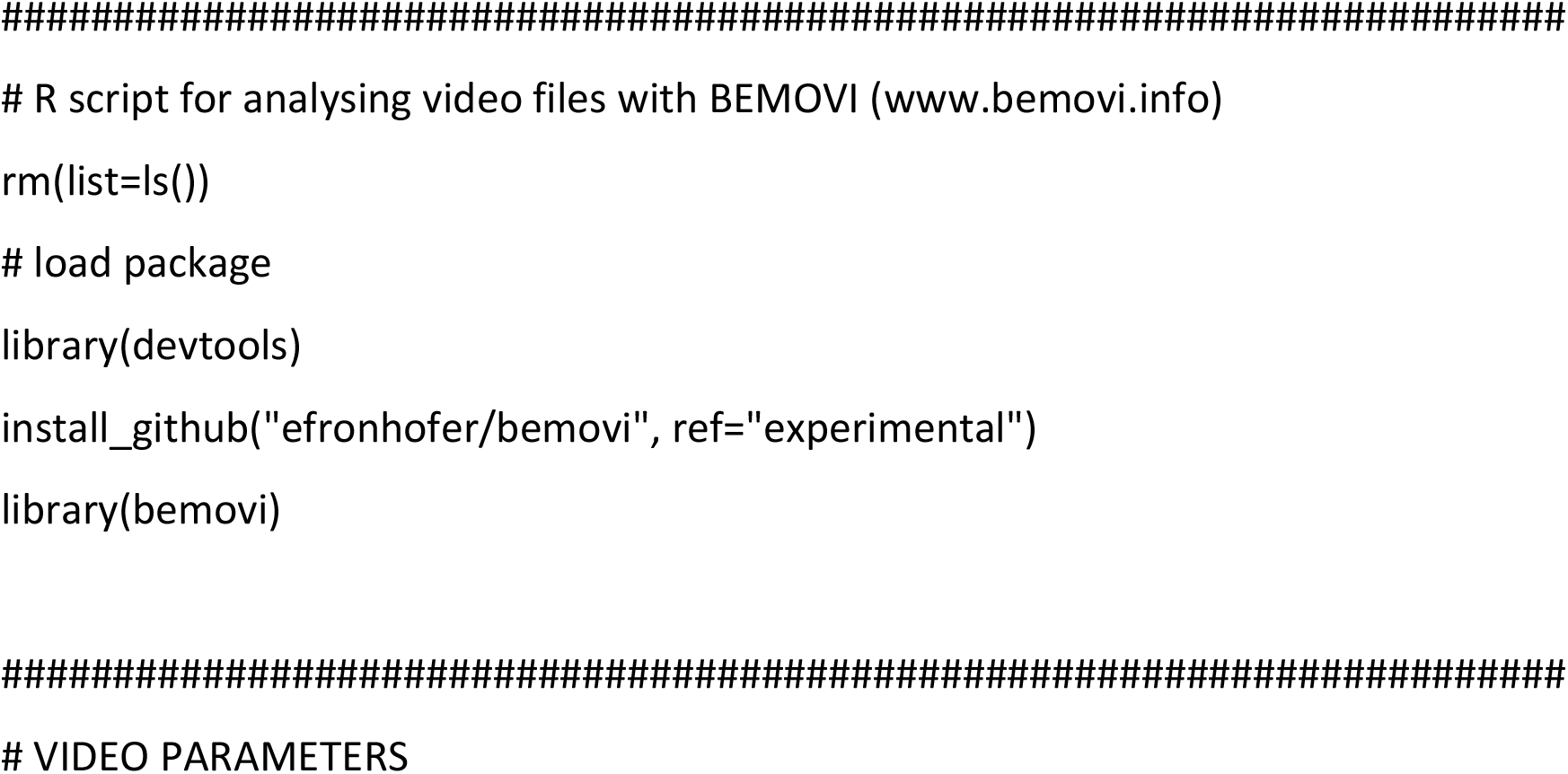
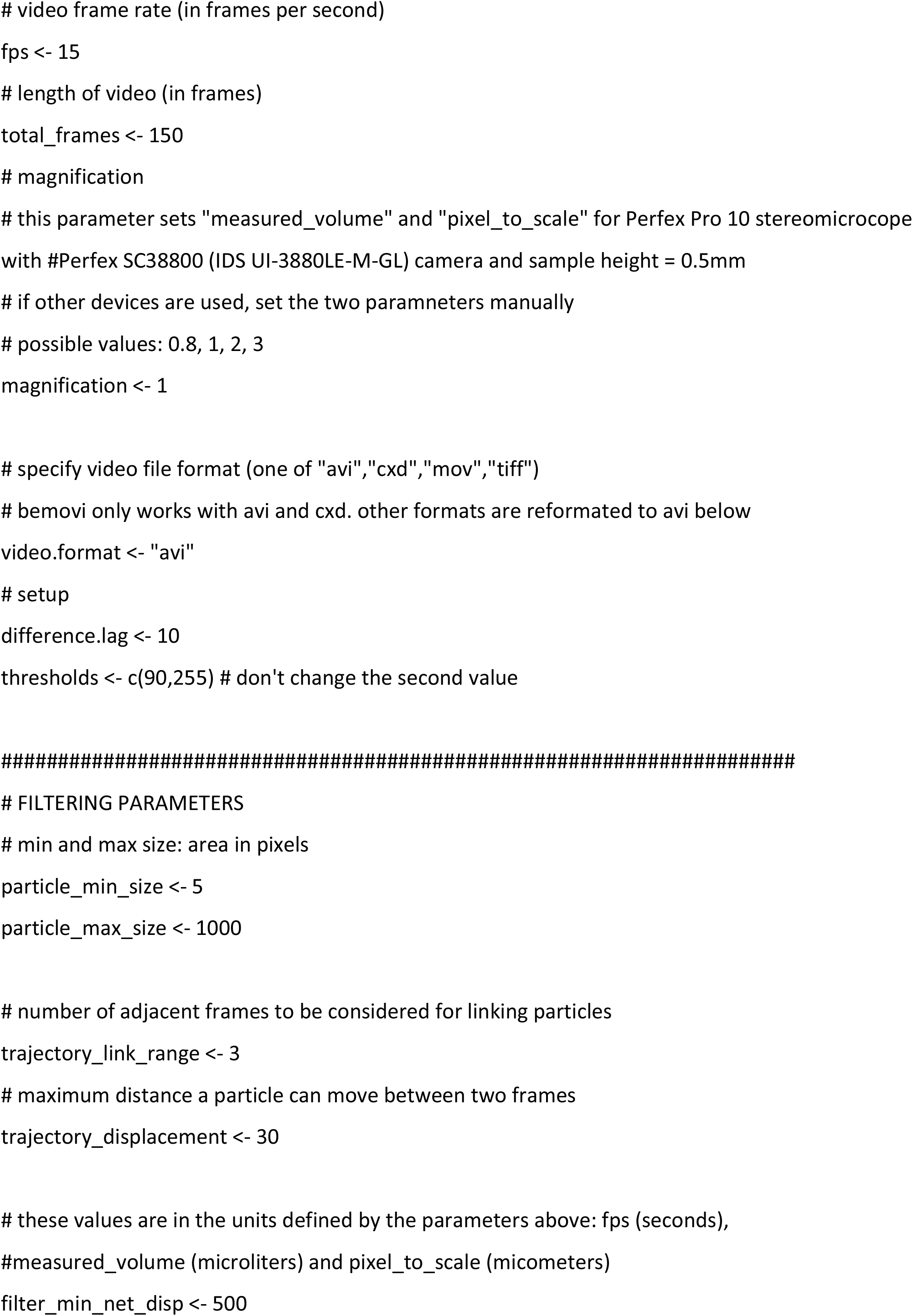
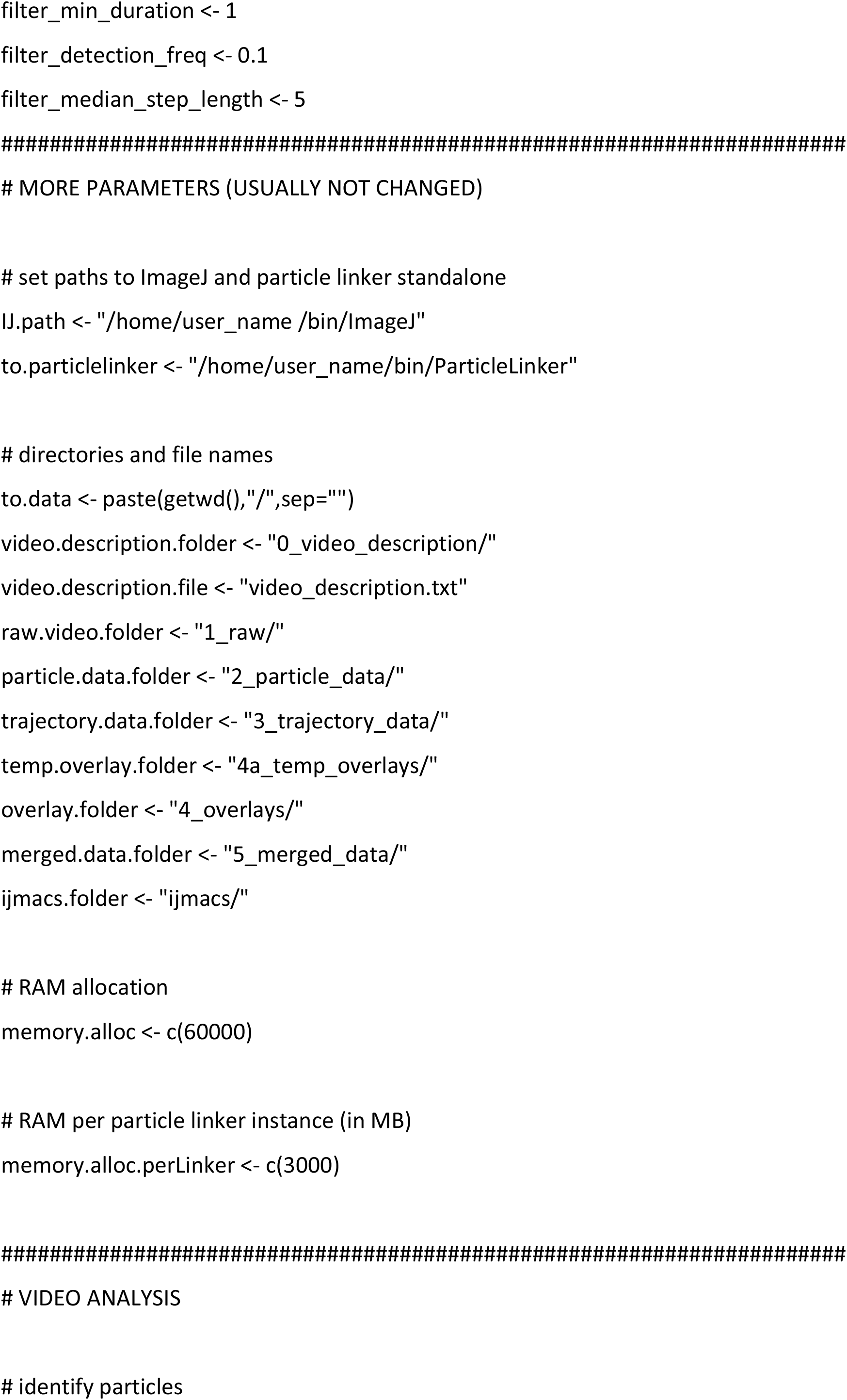
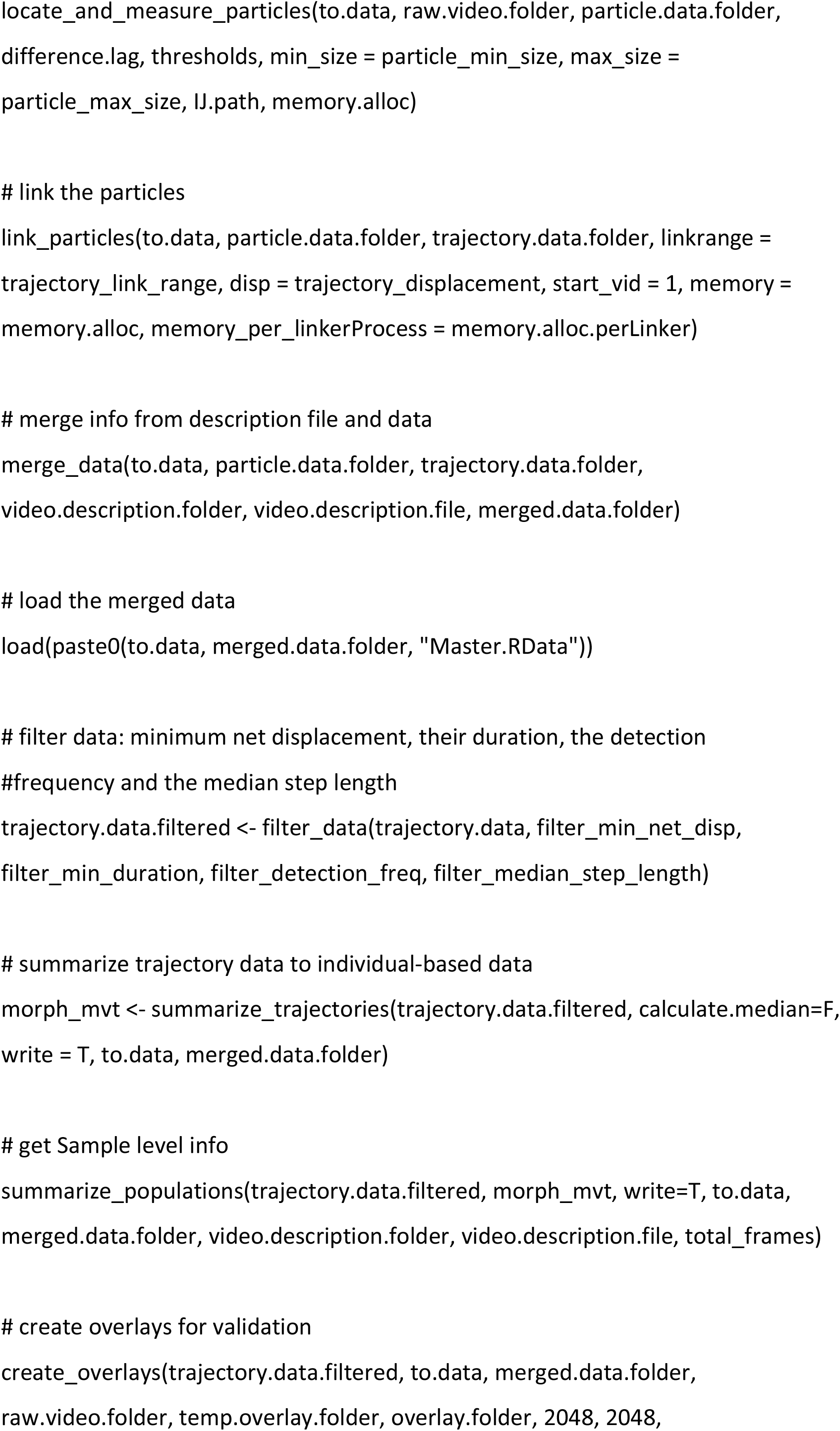
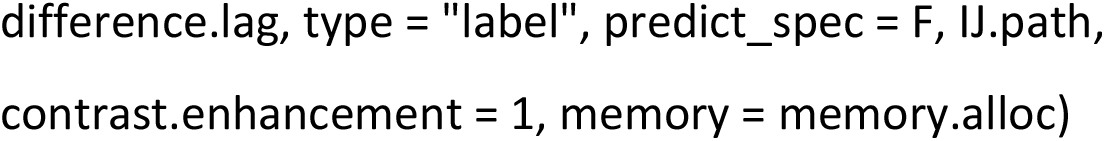

### S3 Trait correlations

**Figure S3.**
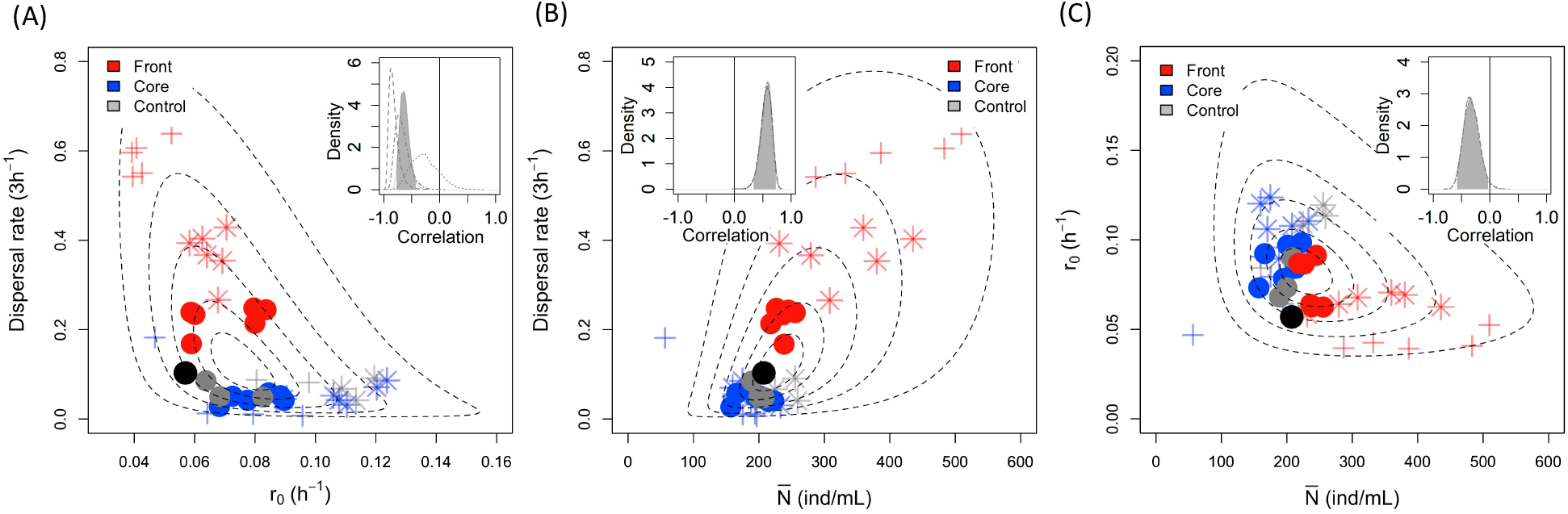
Overall correlation between (A) dispersal - *r0*, (B) dispersal - *N̅* and (C) *r0* - *N̅* obtained with Bayesian inference. Symbols are the average values for a given selection line and year, with blue, red and grey corresponding to core, front and control treatment, respectively. Different symbols refer to the three different years: circle (year 1), star (year 2), cross (year 3). The black point represents the ancestral values (overall mean) of the founder population. The ellipses are bound to non-linear space and correspond to the 10, 25, 50, 75 an 95 % CI of the correlation of pairs of traits. The shaded areas in the insert panels represent the posterior distribution of the overall correlation coefficients (across selection treatments and years). The dot-dashed lines show the posterior distribution of the year 1 correlation coefficients, dashed lines of the year 2, and dotted lines of the year 3. The black line in the inserts highlights the 0 value, and thus the absence of correlation.

### Script correlation analysis

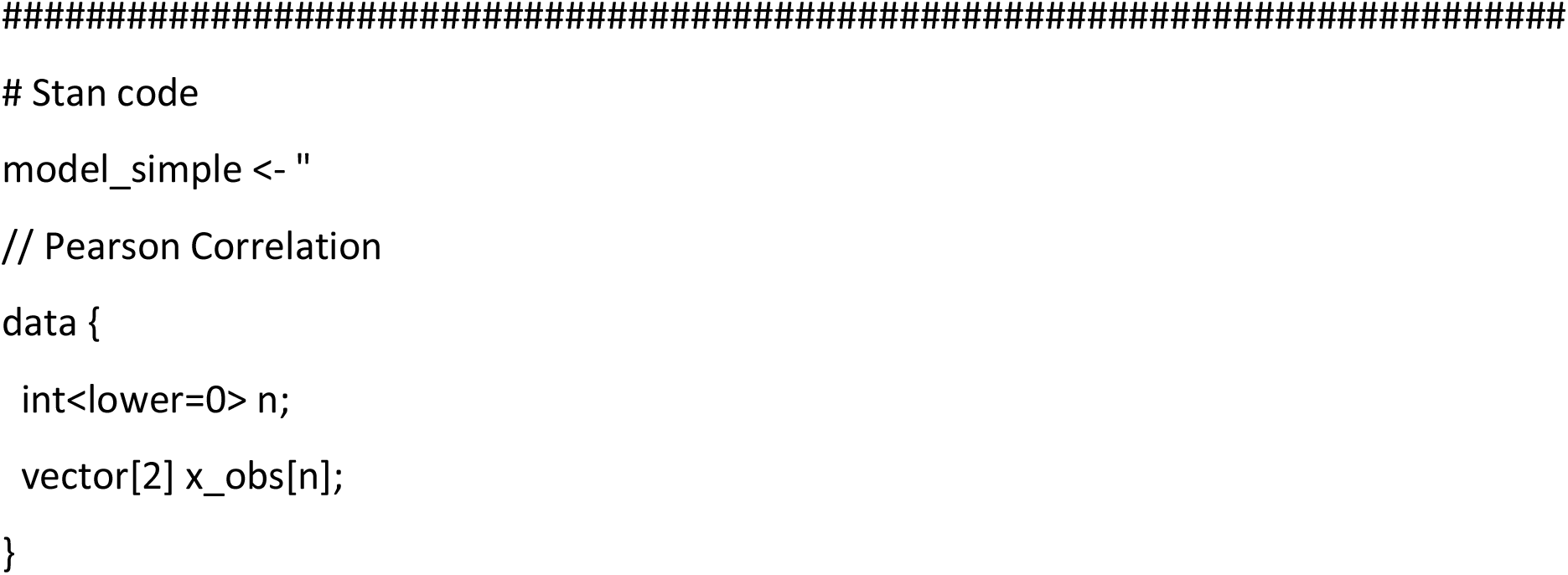
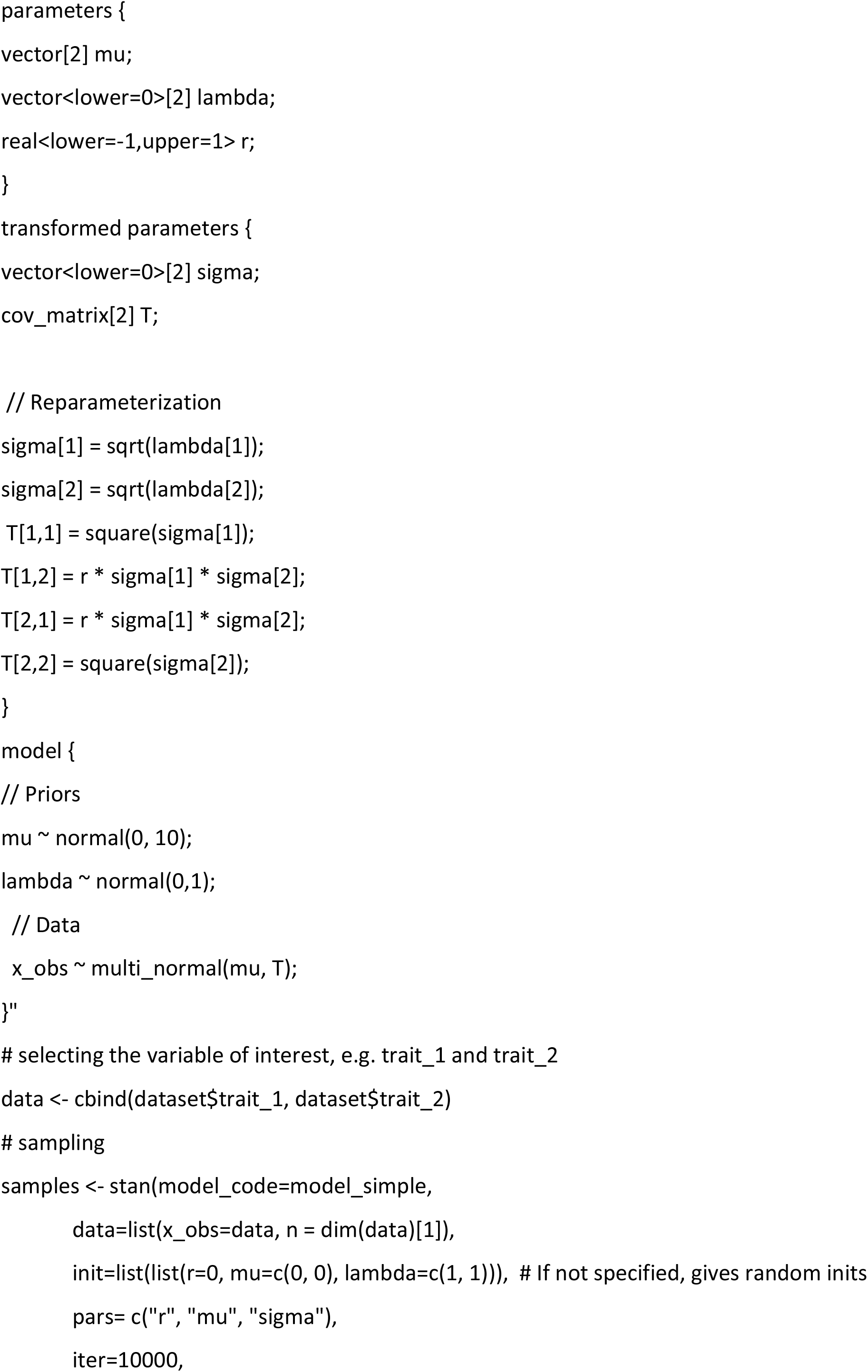
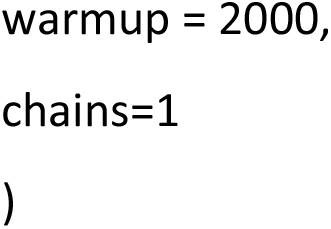

### S4 Founder population trait values and correlations

Prior to the start of the long-term experiment, we characterised the 20 founder strains for dispersal and population growth characteristics (*r_0_*, *N̅*), with 3-4 replicates per strain, as described in the main text. Using a Bayesian approach (Rosenbaum *et al*., 2019), we determined median values and 95% CI for each strain (Table S1, see also section S1). Figure S4 illustrates the (bivariate) trait space occupied by the mix of the strains in the founder population. For example, Fig. S4A shows considerable genotypic variation in both dispersal and *r_0_*. Certain strains have very high *r_0_* and very low dispersal, and several strains have relatively high dispersal and intermediate levels of *r_0_*. As shown in the main text, these two types of strains are targeted by short-term selection in the range core and front treatments, respectively. There are no strains with very high levels of both dispersal and growth, and such variants also do not seem to evolve in the long term (see Fig. 4), suggesting that this part of the trait is unavailable to the genetic backgrounds used in this experiment.

**Figure S4.**
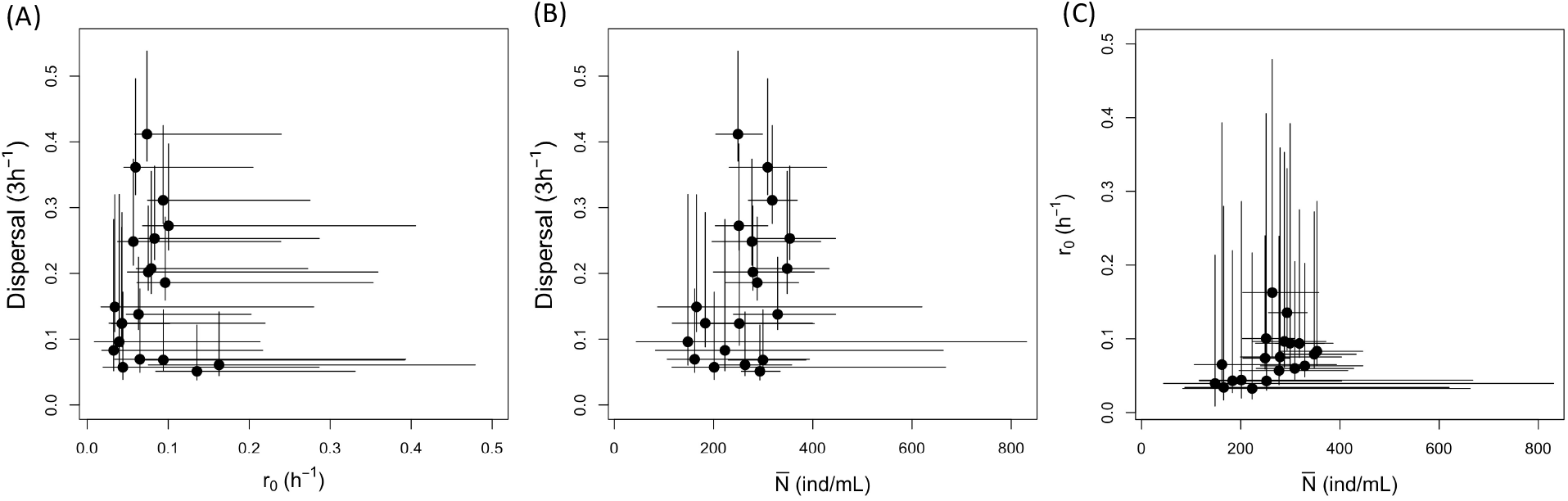
Trait relationships in the base population (mix of the 20 founder strains) for (A) dispersal - *r0*, (B) dispersal - *N̅* and (C) *r0* - *N̅*. Each point represents the median trait values of a strain with the 95% CI (see also Table S1).

### S5 Trait relationships relative to the control treatment

In the main text, Figure 4A-C illustrates short- and long-term trends in pairwise trait associations, in relation to the model predictions. The long-term trends are inferred from the comparison of measurements taken at different time points (year 1, 2, 3, see main text), and we can therefore not a priori exclude the possibility that the (evolutionary) change in a trait is confounded with a measurement year effect. Ideally, to avoid this problem, samples would be frozen each year and all samples measured at the same time in a single assay at the end of the long-term experiment. However, freezing of samples was not possible for our lines. Instead, we accounted for potential year effects by expressing the performance of range core and range front lines relative to the control treatment. This was done by subtracting the means of the control lines from the values of individual core or front lines from the same year. Patterns for these standardized trait associations (Fig. S5) are very similar to those shown in Fig. 4A-C. Thus, our main conclusions regarding the divergence of selection lines were unlikely to be affected by measurement year effects.

**Figure S5.**
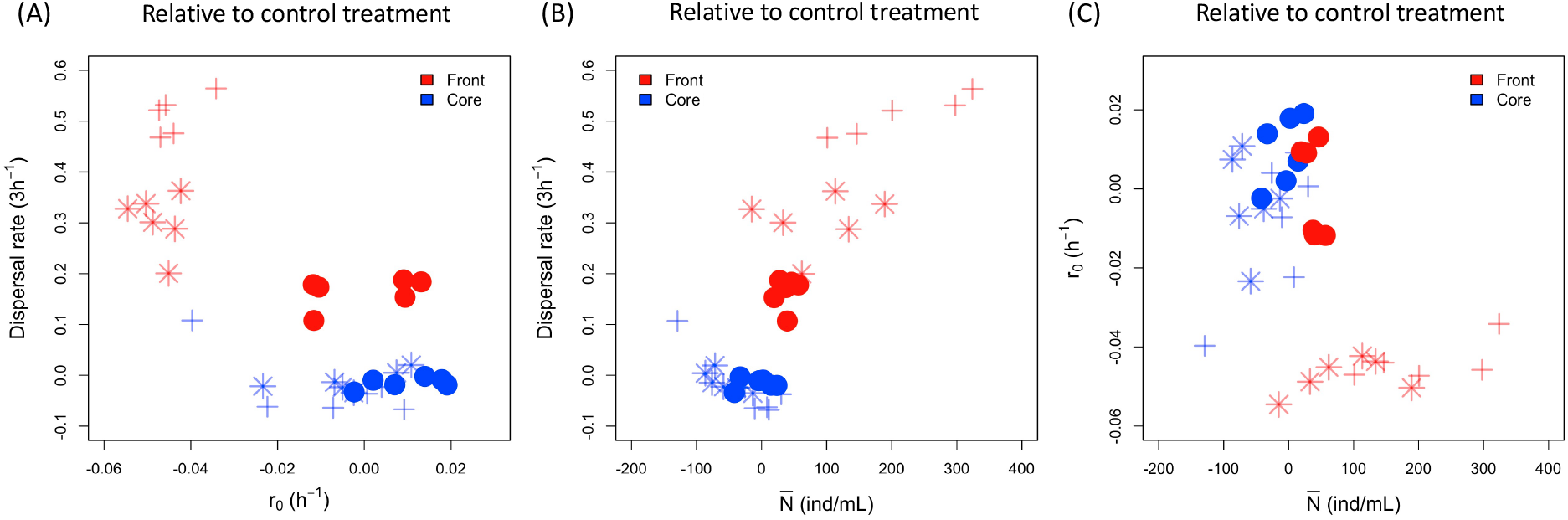
Trait relationships between (A) dispersal - *r0*, (B) dispersal – *N̅* and (C) *r0* – *N̅*, expressed relative to the control treatment (core/front minus control values), for each of three years. Symbols are the average values for each selection line. Negative values correspond to decreased trait values compared to the control treatment of the same year, positive indicated increased values and 0 corresponds to no changes. Different symbols refer to the three different years: circle (year 1), star (year 2), cross (year 3).

### S6 Complementary experiments

After the long-term experiment was completed, additional tests were performed with the evolved selection lines. First, we performed a ‘treatment-reversal’ experiment. To this end, we divided the evolved lines in two new replicates. The first replicate was continued in the original treatment, whereas the second replicate was subjected to the other (opposite) treatment. This experiment was run for 9 cycles and dispersal measured, following the protocols described in the main text.

#### Results

Range front lines continued to show higher dispersal than range core lines, when switched to ‘range core’ selection conditions (Fig. S6.1A). Conversely, range core lines continued show very low dispersal, when switched to ‘range front’ selection conditions (Fig. S6.1B).

**Figure S6.1.**
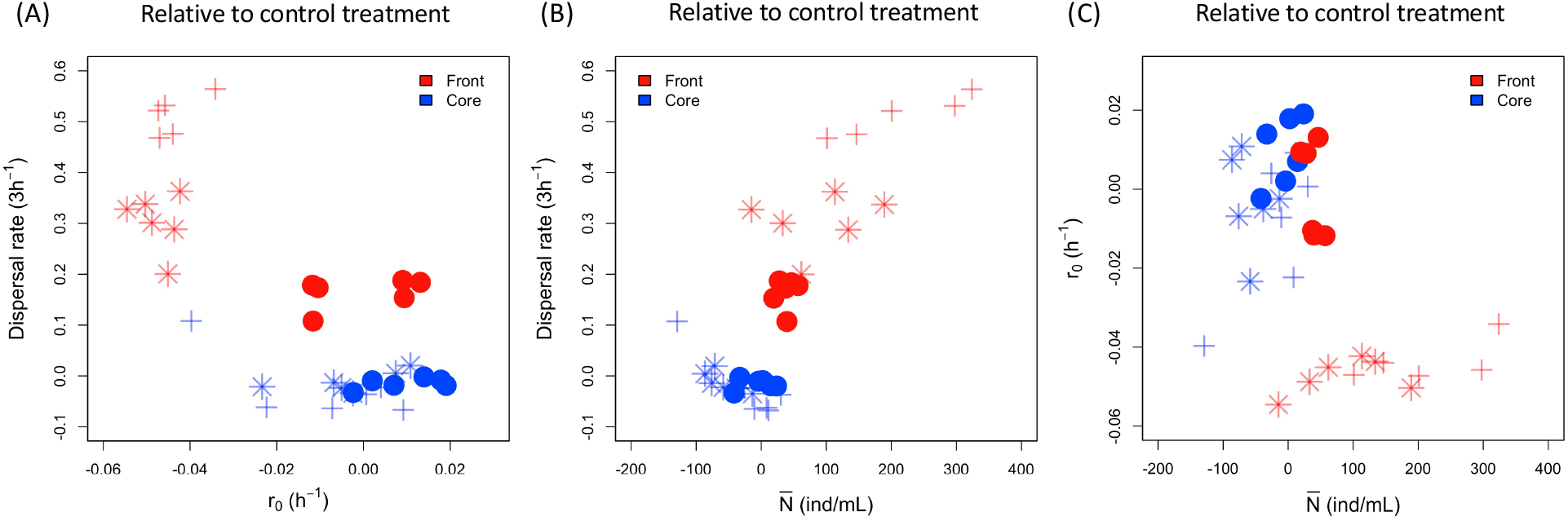
Treatment-reversal experiment. (A) Front and core evolved lines exposed for 9 cycles to core treatment. (B) Front and core evolved lines exposed for 9 cycles to front treatment.

Secondly, we wanted to test whether the observed treatment effects in the long-term experiment, as measured in single-line assays, were strong enough to be picked up by selection. To this end, we mixed range core and front lines at different ‘initial’ proportions, and then exposed these mixes (together with pure 100% core and front controls) to a range core or range front treatment for 3 cycles.

#### Results

Under range core selection (Fig. S6.1A), we find a decrease in dispersal in the mixes, reaching levels as low as those observed for pure range core lines, whereas pure front lines continue to show high dispersal. Conversely, under range front selection (Fig. S6.1B), we find an increase in dispersal in the mixes, reaching levels comparable to values observed for the pure range front lines. Pure range core lines also show an increase in dispersal, but still disperse less than the mixes or the pure front lines. These results indicate a match between selection history and selection treatment, meaning that front lines have a selective advantage under front selection and core lines under core selection. Indeed, in similar experiments (F. Manzi & O. Kaltz, unpublished), we find that such observed phenotypic changes go hand in hand with the fixation of range core or front lines.

**Figure S6.2.**
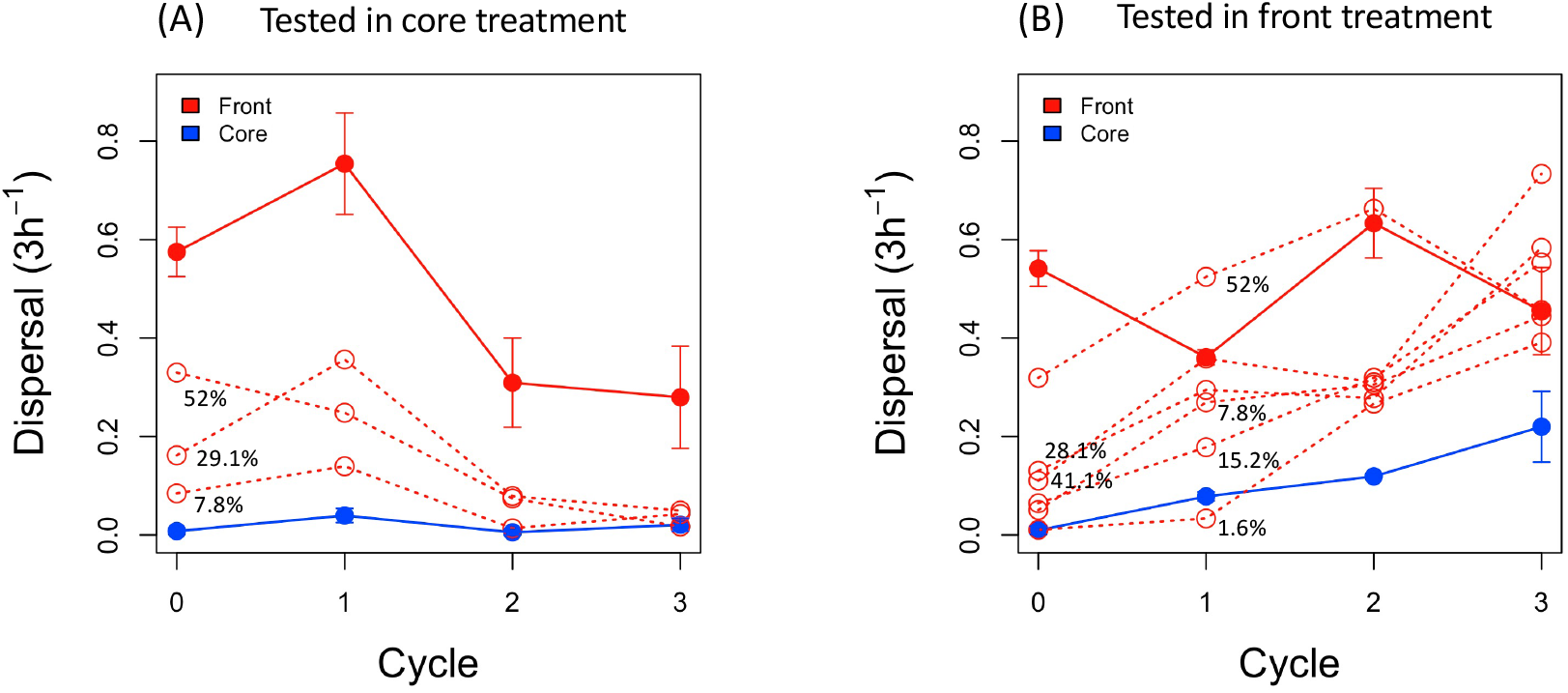
Mixed-lines experiment, with (A) range core selection or (B) range front selection treatment. Solid connecting lines are pure (100%) evolved core and front selection lines, respectively. Dashed red lines are mixes of core and front lines, with the initial proportion of front lines ranging from 1.6% to 52%. Each dashed line represents a single experimental ’mixed’ replicate, the solid ’pure’ lines represent averages (± SE) over 3 experimental replicates.

### S7 Strain winning probability

The model predictions show strong variation in the winning probability among the 20 strains, i.e. the probability of strains fixation in the population for each of three treatments at the end of the experiment. Although the model is deterministic, we parametrize the model with draws from posteriors (section S4 above). Thus, the model takes into account the data uncertainty and gives a distribution of likely outcomes. Details of the model are given in the main text.

**Figure S7.**
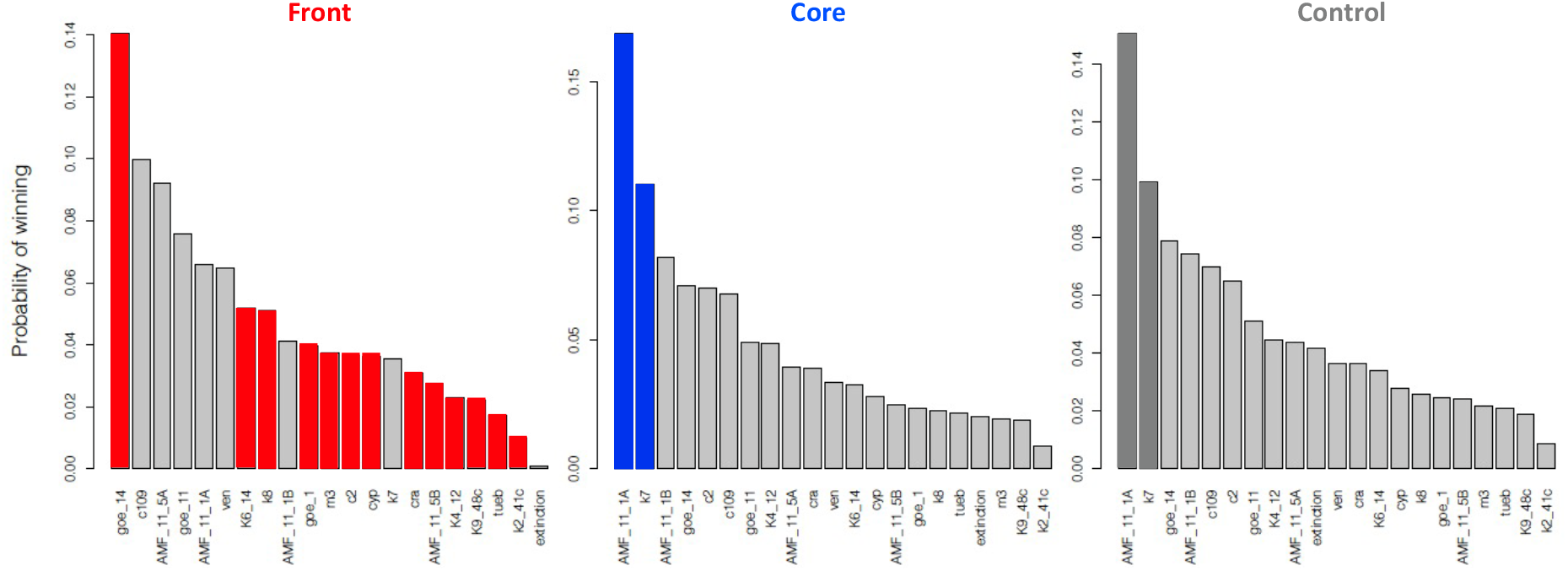
Histograms of strain winning probability for Front, Core and Control treatment with a quasi-extinction threshold of 0.7. Strain winning probability corresponds to the fixation probability among the 20 strains in 10000 model runs. The true potential winner candidates with the right COI genotype are highlighted in red for the Front, blue for the Core and grey for the Control treatment.

### S8 Quasi-extinction threshold

The quasi-extinction threshold implies that strains go extinct if they exhibit densities below this value. The model scenario that fits the observed data indicates a large extinction threshold of 0.7, leading to a similar selection on dispersal and growth rate. Under these conditions, selection favours strains with high dispersal but also a relative high growth rate. When running additional model scenarios with decreased quasi-extinction threshold, selection for growth rate overrides selection for dispersal. Despite the bottlenecks occurring during the dispersal phase, strains with low dispersal can still reach the new patch and regrow to high density. Under these alternative conditions few extinctions occur and all strains can reach the new patch, but it is the strain with the highest growth rate (AMF_11_1A) that becomes fixed in all treatments.

**Figure S8.1.**
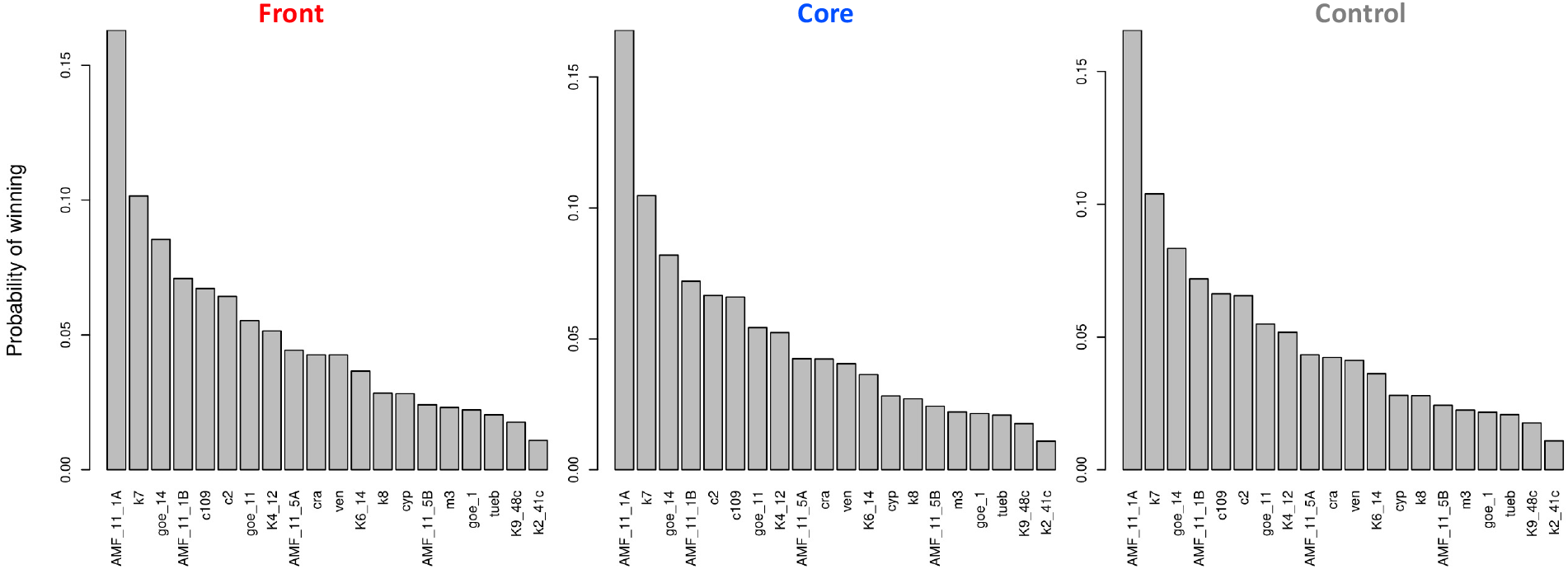
Histograms of strain winning probability for Front, Core and Control treatment with a quasi-extinction threshold 0.001.

**Figure S8.2.**
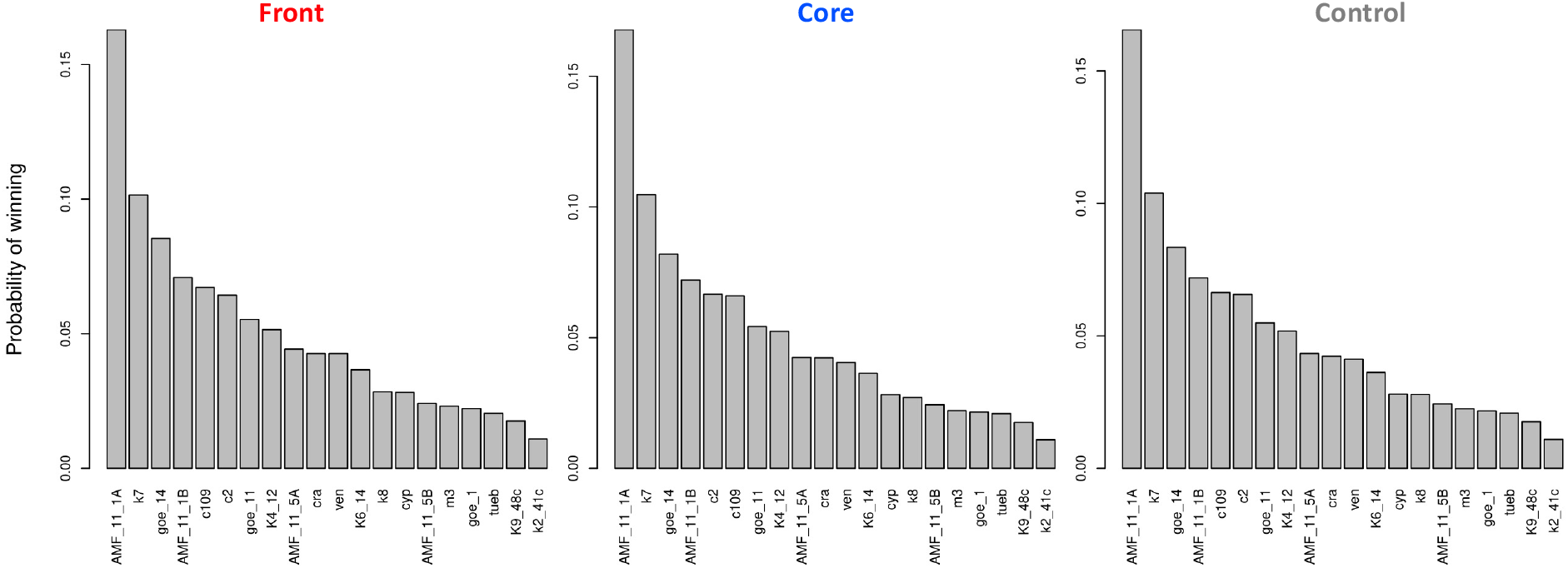
Histograms of strain winning probability for Front, Core and Control treatment with a quasi-extinction threshold 0.1.

**Figure S8.3.**
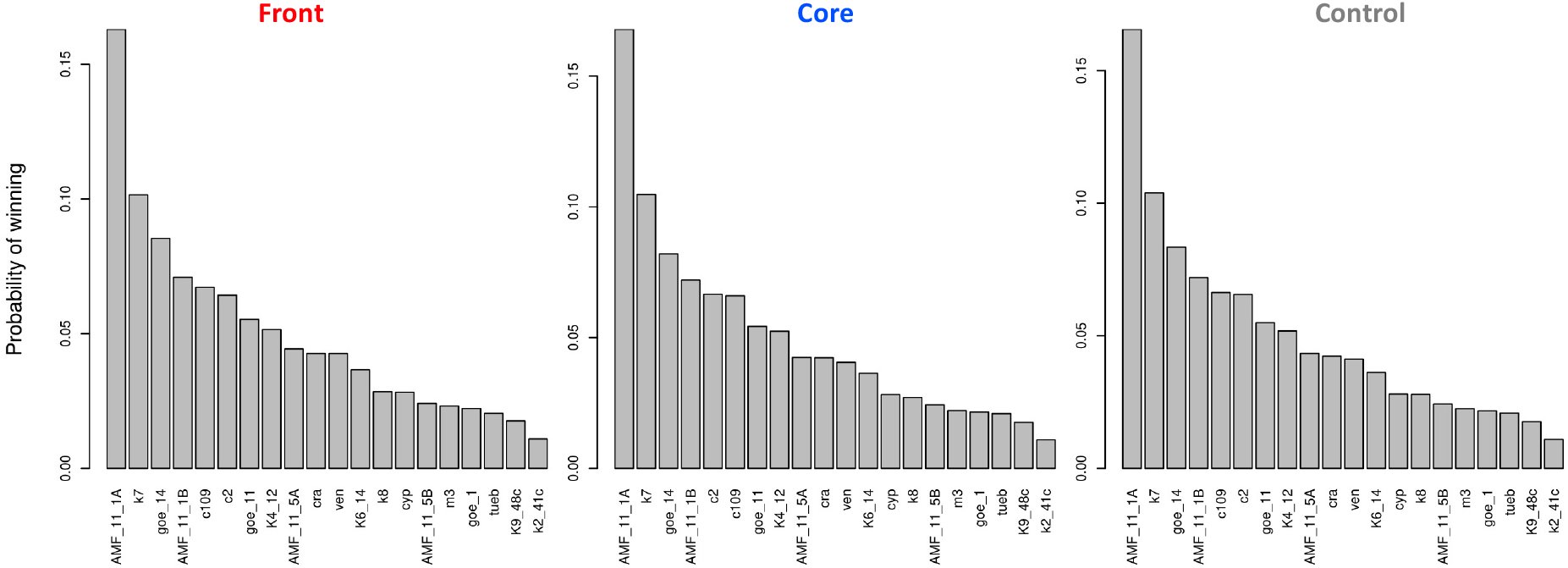
Histograms of strain winning probability for Front, Core and Control treatment with a quasi-extinction threshold 0.5.

**Table S1.** Details and trait values for each of the 20 strains of the founder population.

| Strain | Origin (provided by) | In OK lab since | COI genotype | COI genotype detected at cycle 30 | Median dispersal | 95% CI dispersal | Median r0 | 95% CI r0 | Median $\bar{N}$ | 95% CI $\bar{N}$ |
| --- | --- | --- | --- | --- | --- | --- | --- | --- | --- | --- |
| AMF_11_1A | Russia (A. Potekhin) | 2014 | b05 | range core and control | 0.060 | 0.044; 0.141 | 0.162 | 0.075; 0.479 | 263.5 | 204.0; 357.4 |
| AMF_11_1B | Russia (A. Potekhin) | 2014 | b04 | not detected | 0.068 | 0.052; 0.144 | 0.093 | 0.063; 0.391 | 299.5 | 229.7; 386.2 |
| AMF_11_5A | Russia (A. Potekhin) | 2014 | * | not detected | 0.311 | 0.275; 0.424 | 0.093 | 0.074; 0.275 | 318.3 | 270.1; 368.4 |
| AMF_11_5B | Russia (A. Potekhin) | 2014 | b07 | range front | 0.124 | 0.088; 0.292 | 0.042 | 0.026; 0.219 | 183.3 | 116.7; 399.1 |
| c109 | Unknown | 2014 | b01 | not detected | 0.185 | 0.159; 0.285 | 0.096 | 0.061; 0.353 | 288.0 | 223.3; 371.7 |
| c2 | Isolated from commercial mix | 2004 | b07 | range front | 0.069 | 0.049; 0.176 | 0.065 | 0.033; 0.393 | 162.0 | 106.5; 393.3 |
| cra | Kraków, Poland | 2006 | b07 | range front | 0.095 | 0.060; 0.320 | 0.039 | 0.008; 0.213 | 148.2 | 44.2; 830.9 |
| cyp | Cyprus | 2003 | b07 | range front | 0.148 | 0.111; 0.319 | 0.034 | 0.016; 0.279 | 165.6 | 87.3; 620.0 |
| goe_1 | Stuttgart, Germany | 2008 | a18** | not detected | 0.248 | 0.212; 0.373 | 0.056 | 0.037; 0.239 | 277.3 | 196.7; 415.2 |
| goe_11 | Stuttgart, Germany | 2007 | b01 | not detected | 0.202 | 0.174; 0.302 | 0.075 | 0.049; 0.358 | 279.3 | 199.5; 403.2 |
| goe_14 | Stuttgart, Germany | 2007 | b07 | range front | 0.272 | 0.235; 0.397 | 0.100 | 0.068; 0.405 | 251.0 | 203.7; 309.2 |
| k2_41c | Japan, parents KNZ 5 & KNZ 2 | 2001 | b07 | range front | 0.123 | 0.090; 0.270 | 0.042 | 0.030; 0.102 | 251.9 | 181.6; 403.3 |
| K4_12 | Japan, parents KNZ 5 & KNZ 2 | 2001 | b07 | range front | 0.057 | 0.038; 0.171 | 0.044 | 0.019; 0.286 | 201.3 | 116.6; 667.8 |
| K6_14 | Japan, parents KNZ 5 & KNZ 2 | 2001 | b07 | range front | 0.207 | 0.169; 0.355 | 0.079 | 0.060; 0.272 | 348.3 | 273.9; 432.9 |
| k7 | Japan, parents KNZ 5 & KNZ 2 | 2001 | b05 | range core and control | 0.051 | 0.037; 0.121 | 0.135 | 0.084; 0.330 | 293.4 | 256.0; 334.2 |
| k8 | Japan, parents KNZ 5 & KNZ 2 | 2001 | b07 | range front | 0.411 | 0.370; 0.537 | 0.073 | 0.058; 0.239 | 249.1 | 204.4; 298.5 |
| K9_48c | Japan, parents KNZ 5 & KNZ 2 | 2001 | b07 | range front | 0.137 | 0.114; 0.224 | 0.063 | 0.048; 0.202 | 329.1 | 239.9; 446.2 |
| m3 | Isolated from commercial mix | 2004 | b07 | range front | 0.361 | 0.319; 0.495 | 0.059 | 0.045; 0.204 | 309.0 | 231.1; 427.8 |
| tueb | Tübingen, Germany (H-D Görtz) | 2001 | b07 | range front | 0.082 | 0.051; 0.282 | 0.032 | 0.018; 0.216 | 223.0 | 82.9; 662.9 |
| ven | Venice, Italy | 2006 | a01** | not detected | 0.253 | 0.220; 0.363 | 0.083 | 0.063; 0.286 | 353.7 | 276.8; 446.0 |

\* identified as *Paramecium multinucleatum*
\*\* determined in Killeen, J., Gougat-Barbera, C., Krenek, S. & Kaltz, O. (2017). Evolutionary rescue and local adaptation under different rates of temperature increase: a combined analysis of changes in phenotype expression and genotype frequency in *Paramecium* microcosms. *Molecular Ecology*, 26, 1734-1746.

